# The biophysical mechanism of mitochondrial pearling

**DOI:** 10.1101/2024.12.21.629509

**Authors:** Gabriel Sturm, Kayley Hake, Austin E.Y.T. Lefebvre, Caleb J. Rux, Daria Ivanova, Alfred Millett-Sikking, Kevin M. Tharp, Beiduo Rao, Michael Closser, Adam James Waite, Magdalena Precido-Lopez, Alex T Ritter, Sophie Dumont, Wen Lu, Suliana Manley, Juan C. Landoni, Wallace F. Marshall

## Abstract

Mitochondrial networks exhibit remarkable dynamics that are driven in part by fission and fusion events. However, there are other reorganizations of the network that do not involve fission and fusion. One such exception is the elusive, “beads-on-a-string” morphological transition of mitochondria. During such transitions, the cylindrical tubes of the mitochondrial membrane transiently undergo shape changes to a string of “pearls” connected along thin tubes. These dynamics have been anecdotally observed in many contexts and given disparate explanations. Here we unify these observations by proposing a common underlying mechanism based on the biophysical properties of tubular fluid membranes for which it is known that, under particular regimes of tension and pressure, membranes reach an instability and undergo a shape transition to a string of connected pearls. First, we use high-speed light-sheet microscopy to show that transient, short-lived pearling events occur spontaneously in the mitochondrial network in every cell type we have examined, including primary fibroblasts, T-cells, neurons, and budding yeast. We present evidence that transient mitochondrial pearling occurs during important biological events, particularly during T cell activation, neuronal firing, and replicative senescence. Using our high-temporal resolution data, we identify two distinct categories of spontaneous pearling, i) internal pressure-driven pearling generated by ionic flux, and ii) external tension-driven pearling generated by the cytoskeleton. By applying live-cell STED and FIB-SEM imaging we document the structural reorganization of inner cristae membranes during mitochondrial pearling and the role of the MICOS complex in regulating the frequency of pearling events. We then establish numerous methods for inducing pearling, including the ability to induce these dynamics with single mitochondrion precision. These methods include ionophores, channel activators, osmotic shock, detergents, laser stimulation, membrane intercalating molecules, chemical fixation, and micro-needle force. These disparate inducers establish three main physical causes of pearling, i) ionic flux producing internal osmotic pressure, ii) membrane packing lowering bending elasticity, and iii) external mechanical force increasing membrane tension. Pearling dynamics thereby reveal a fundamental biophysical facet of mitochondrial biology. We suggest that pearling should take its place beside fission and fusion as a key process of mitochondrial dynamics, with implications for physiology, disease, and aging.

## Introduction

Mitochondria form networks that continually reorganize via fission and fusion (1, 2). A less appreciated behavior of mitochondria is the transient formation of “beads-on-a-string” morphology (**Fig. 1A**), first observed in 1915 by Margaret and Warren Lewis (3). This morphology has been observed in numerous biological contexts and has been referred to as “beads-on-a-string” (4), “COMIC” (5), “MOAS” (6), “nodules”, “condensation”, “beading”, “fragmentation sites” (4, 7), and “thread-grain transitions” (8). Sightings include the muscle cells of *C. elegans* (9), the hypocotyl cells of Arabidopsis plants (10), during the growth phase and meiosis of budding yeast (11, 12) and mitochondrial disease patient-derived cells (13). Additionally, beads-on-a-string mitochondria have been observed in postmortem brains of healthy, aged, Alzheimer’s, and hypoxic-treated mice and humans (6, 14, 15).

**Figure 1.**
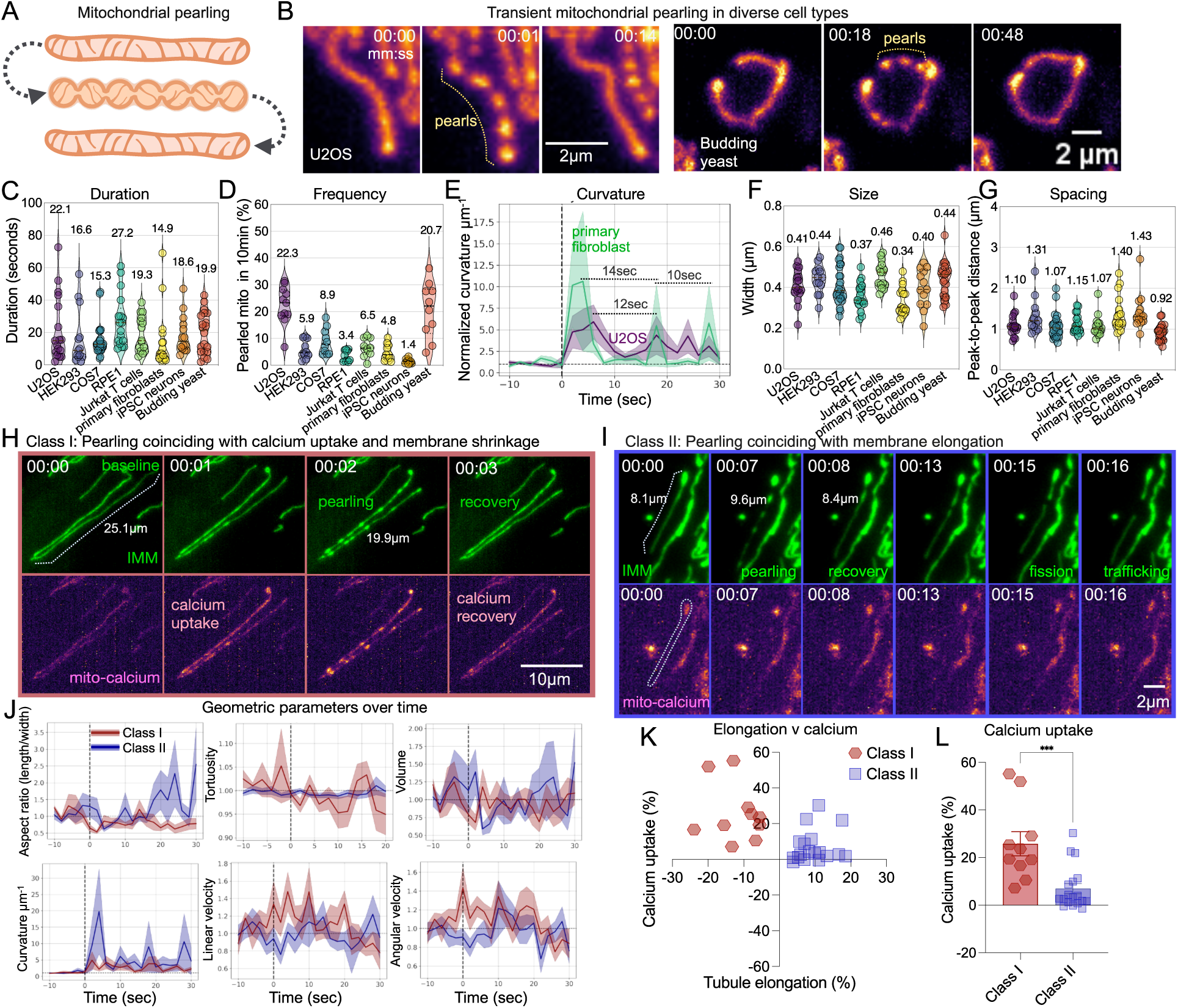
Spontaneous pearling is conserved and encompasses two distinct biophysical events. (**A**) Schematic of a transient mitochondrial pearling event. (**B**) Representative examples of transient pearling along the inner mitochondrial membrane of U2OS (COX8a-mEmerald), Jurkat T cells (COX8a-StayGold) and budding yeast (pre-COX4-mNeonGreen) (**C**) Duration of pearling transitions from uniform to pearl back to uniform transition amongst 8 different cell types (n= 13-25 events per cell type). (**D**) Rate of pearling events measured as the probability any mitochondria will pearl over a 10 minute period (n=10-11 videos per group). (**E**) Normalized curvature per µm across the length of mitochondrial tubule over time for primary fibroblasts (n=19 events) and U2OS cells (n=18 events). Each event is normalized to the average of the 3 initial timepoints. Dotted vertical line indicates the onset of pearling. (**F-G**) Mean cross-diameter width (**F**) and distance between the center of each neighboring pearl (**G**) for listed cell types (n=13-25 events per cell type). (**H-I**) Representative time series of tubule (G) shrinking and (H) elongation during a spontaneous pearling event in primary human fibroblasts co-labeled with the mitochondrial inner-membrane (COX8a-mStayGold) and mitochondrial calcium (genetically-encoded mito-R-GECO1). Dotted line indicates length of adjacent mitochondrial tubule. (**J**) Structural and dynamical metrics comparing pearls that undergo stretching (red) and shrinking (blue). All values normalized the average of the first three timepoints. Dotted line indicates the onset of pearling. Error bars = S.E.M. (**K**) Calcium uptake vs tubule elongation amongst spontaneous pearling events. Blue circles and red dots mark events under which elongation or shrinking occurred, respectively. (**L**) Comparison of calcium uptake between elongated and shrunk mitochondria during pearling. ** = p < 0.01 performed by non-parametric Mann-Whitney test. n = 10-20 events per group.

It has been proposed that the mitochondrial fission machinery based on dynamin-like protein Drp1, could provide a force to constrict the mitochondrial membrane into pearls. However, DRP1 knock-out cells have increased rather than decreased pearling rates (5, 16). Other hypothesized explanations for this phenomenon include reactive oxygen species (ROS) generation (17), ER constrictions around the mitochondrial tube, cytoskeleton remodeling, and non-conventional permeability transition pores (mPTP, (18)), but none have been able to fully explain the experimental observations. More recently, the beads-on-a-string morphology was explained by ER contact-sites where calcium influxes into the mitochondrion (19). But pearl formation can be independent of calcium influx (15), (20). Thus, the mechanism by which pearls form is unclear.

A different mechanism for these dynamics comes from the physics of the Rayleigh-Plateau instability governed by the Helfrich model of elastic membranes (21–23). A cylinder is not actually the minimal surface area shape for an elongated membrane tube. If the surface is folded into sinusoidal curves, the same internal volume can be enclosed with a smaller surface area. Membrane tubes tend to be cylindrical because the energy of bending a membrane is substantially larger than the energy of stretching it, thus the lowest energy state is one in which the surface is minimally curved (cylinder) at the expense of having to stretch to accommodate the internal volume. If membrane tension is increased, by inflating the internal volume or by applying mechanical pulling forces, the energy balance shifts such that the stretching term becomes larger than the bending term, resulting in a lower energy state in which the membrane curves into pearls (23–26).

Any cylindrical tube of membranes, including double membranes such as cellular membranes, and reticulated organelle networks like mitochondria will undergo a spontaneous shape transition when the energy stored by membrane stretching exceeds the elastic energy stored by membrane bending. These energies are determined by the organelle’s physical properties; in particular, membrane tension and elastic modulus. The membrane can reach this threshold from i) increased internal osmotic pressure, ii) increased external mechanical tension, or iii) reduced bending elasticity (i.e. stiffness) (23, 27). In accordance with this mechanism, we refer to this dynamical phenomenon as mitochondrial pearling.

Such pearling-like forms have been described in plasma membranes (28, 29), axons (30, 31), microtubule-associated proteins (32), contractile vacuoles (33) and the endoplasmic reticulum (ER) (34–36). However, because the mitochondrion has an inner membrane that is electrochemically coupled and highly structured into helicoid cristae (37), it has an elastic modulus an order of magnitude greater than typical double membranes of its size (21). These properties make it particularly resistant to deformation and thereby physically unique among cellular organelles.

Here we propose the Rayleigh-Plateau mechanism may provide a unified model of mitochondrial pearling that can account for the many circumstances in which it has been observed. Using live-cell imaging, high-resolution structural information, and perturbation experiments, we find that increased osmotic pressure or mechanical tension, or altering the mitochondrial membrane, can induce a similar pearling instability. The ability of such disparate causes to drive mitochondrial pearling can most parsimoniously be explained if they all work by altering the balance of tension and bending energies in the membrane, thus avoiding the need to invoke a collection of distinct molecular processes.

## Results

### Spontaneous mitochondrial pearling encompasses two physically distinct events

Live-cell time lapse imaging of individual mitochondria tubules revealed that mitochondria will occasionally transition into pearled units connected by thin membrane tubules, which persist transiently before returning to a uniform shape (**Fig. 1B; Movie S1**). These events have previously been observed in standard imaging conditions in mammalian cells such as U2OS, HEK293, HepG2, MEFs, and cortical neurons (5, 15, 16, 20). Those reports documented events lasting 5 seconds to 2 minutes, with each mitochondrion having a 10%-50% chance of pearling every 10 minutes, depending on cell type. Here, we start by extending these reports using a single-objective light-sheet (SOLS) microscope (a.k.a. “Snouty”, (38)) to image whole-cell mitochondrial networks in 3D at 2 Hz (i.e. 2 volumes/second) with minimal phototoxicity compared to standard 3D imaging. We found transient pearling events in every cell type studied, which included U2OS, HEK293, COS7 (monkey), RPE1, Jurkat cells, primary human CD8 T cells, primary human fibroblasts, iPSC-derived neurons, and budding yeast (**Fig. 1B; SI Appendix, Fig. S1A-H; Movie S2, Table S1;**). Using the SOLS microscope, full cylinder-pearl-cylinder transitions occurred in as little as 2 seconds (0.5 Hz) **(Fig. 1C; Movie S3**). Based on these dynamics, a typical confocal microscope (<0.2 Hz) would miss up to 27% of pearling events, depending on cell type. Additionally, imaging with SOLS microscopy at 1 Hz revealed that the probability of a pearling event can vary widely between cell types: In a ten-minute interval, it can be as low as 1.4% of mitochondria in iPSC-derived neurons or as high as 22.3% of mitochondria in U2OS cells (**Fig. 1D; SI Appendix Fig. S1I**). Mitochondria that have pearled tend to undergo 2nd and 3rd events within regular time intervals of 10-14 seconds (**Fig. 1E**). The shape of these pearls across cell types was highly conserved with an average width (i.e cross-diameter) of 320-480 nm and peak-to-peak distance between pearls of 0.8-1.3 µm (**Fig. 1F-G**).

We also observed pearling events during reconstituted physiological events. First, we cultured iPSC-derived neurons and observed spontaneous pearling of individual mitochondrial tubules temporally correlated with baseline action potentials tracked by cytoplasmic calcium flickering (**SI Appendix, Fig. S1E; SI Appendix, Fig. S2A; Movie SS4**), as well as cell-wide mitochondrial pearling during chemically-induced (50 mM KCl) action potentials (**SI Appendix, Fig. S2B; Movie S5**). Second, frequent transient pearls formed at the immune synapses of both primary human CD8 T cells and Jurkat cells when conjugated with Raji cells pulsed with the superantigen Staphylococcus enterotoxin E (SEE) (39) (**SI Appendix, Fig. S2C-E; Movie SS6-8**). Third, primary senescent human fibroblasts serially passaged until replicative senescence(40) (**SI Appendix, Fig. S2F; Movie S9**) had pearling events propagating across the entirety of their hyper-fused mitochondrial networks. These vastly different biological contexts support the ubiquitous and physiological nature of mitochondrial pearling.

Further resolving this temporal transition revealed a wide range of intermediate pearl shapes shared across many cell types (**SI Appendix, Fig. S1A-H; Movie S2**), in some cases with different temporal dynamics. For example, in U2OS cells treated with the electron transport chain (ETC) complex III inhibitor rotenone (5µM for 30min), spontaneous pearling was observed across a 25 µm long tubule (**SI Appendix, Fig. S1J; Movie S10**). One minute later, the same tubule underwent a second transition to the pearled state. However, in this second event an additional fused 20 µm long tubule, that was cylindrical during the first pearling event, synchronously pearled with the original tubule, suggesting that pearling can propagate across physically coupled mitochondrial units.

Co-labeling the mitochondrial inner membrane and mitochondrial calcium (Mito-R-GECO1, (41)) revealed that spontaneous pearling events fall into two distinct classes occurring at similar frequencies. Class I consisted of those preceded by a rise in calcium accompanied by shrinking of the tubule length (**Fig. 1H**). Class II events were those not preceded by a detectable increase in calcium but were accompanied by elongation of the tubule (**Fig. 1I**). Close examination of Class II elongation events revealed the tips of the mitochondria appeared to be pulled into thin protrusions, 180-300 nm in width, projecting outwards 1-6 µm (**Movie S11**), and tended to precede fission events, consistent with motor-driven pulling of mitochondria (5). On the other hand, class I calcium-associated pearling events tended to shrink the mitochondrial length, such that all peripheral branches were compacted into a smaller area (**Movie S3&12**). Tracking structure and movement over time (**Fig. 1J**), showed class II elongation-associated pearling events recovered faster and maintained a straight path throughout the event. A small subset of events displayed both elongation and calcium uptake, but shrinking never occurred without calcium uptake (**Fig. 1K-L**). The ratio of elongation-associated to calcium-associated event frequencies differed by cell type with the highest ratio in primary fibroblasts (8:2), followed by U2OS (7:4), and HEK (1:5) cells. In terms of the Rayleigh-Plateau instability, we hypothesized that these two classes of spontaneous events may reflect two different physical forces, with class I driven by changes to internal ionic pressure that swelled the internal volume and class II driven by external mechanical pulling that stretched the membrane. In both cases the energetic contribution of membrane tension increased until it exceeded the energetic contribution of bending elasticity, at which point the membrane transitioned to the pearled shape to minimize its surface area.

Osmotic pressure drives chemically induced mitochondrial pearling

We next investigated the mechanism of class I pearling events. The most likely source of internal pressure would be uptake of ions into the mitochondrial interior driving water molecules into the mitochondrial matrix creating osmotic pressure. Osmotic shock is known to cause pressure-driven pearling instability in membrane tubes (33). To confirm that internal pressure can pearl mitochondria, we replicated the osmotic shock experiment of Margaret Lewis conducted in 1915 using repetitive media changes between hyper and hypo-osmotic media. By replacing the cell medium (280 mOsm, isotonic) with a hypotonic solution of 0.1x PBS (28 mOsm i.e. low-salt water) pearling was induced across the entire mitochondrial network (**Movie S13**). This was subsequently reversed by swapping media back to the original medium or a balanced salt solution of 1x PBS (280 mOsm) (**Fig. 2A-B; SI Appendix, Fig. S3A**). Additionally, swapping media with hypertonic solution of 10x PBS (2800 mOsm) resulted in mitochondria becoming flaccid spaghetti-like cylinders with minimal movement (**Movie S14**). When the hyper-osmotic media was replaced with hypo-osmotic media, pearls appeared. Repetition between these two conditions converted the mitochondria between flaccid (hypertonic) to pearled (hypotonic) states up to five times until the cells detached from the cover glass (**SI Appendix, Fig. S3B; Movie S14**). A 20 second burst of hyperosmotic shock, followed by a return to isotonic media did not return mitochondria to baseline morphology, but instead resulted in cell wide pearling without a rise in mitochondrial calcium (**Movie S15**). Additionally, hypotonic induced pearling occurred even if mitochondria were depolarized with the proton ionophore FCCP, though depolarized mitochondria did not fully recover to baseline curvature after isotonic rescue (**SI Appendix, Fig. S3C**). Under hypotonic shock, pearling was accompanied by shrinking of the tubule length and calcium influx, which were reversed by a return to an isotonic environment (**SI Appendix, Fig. S3D-F**). Class I spontaneous pearling events may thus share a common mechanism with pearling induced by osmotic pressure.

**Figure 2.**
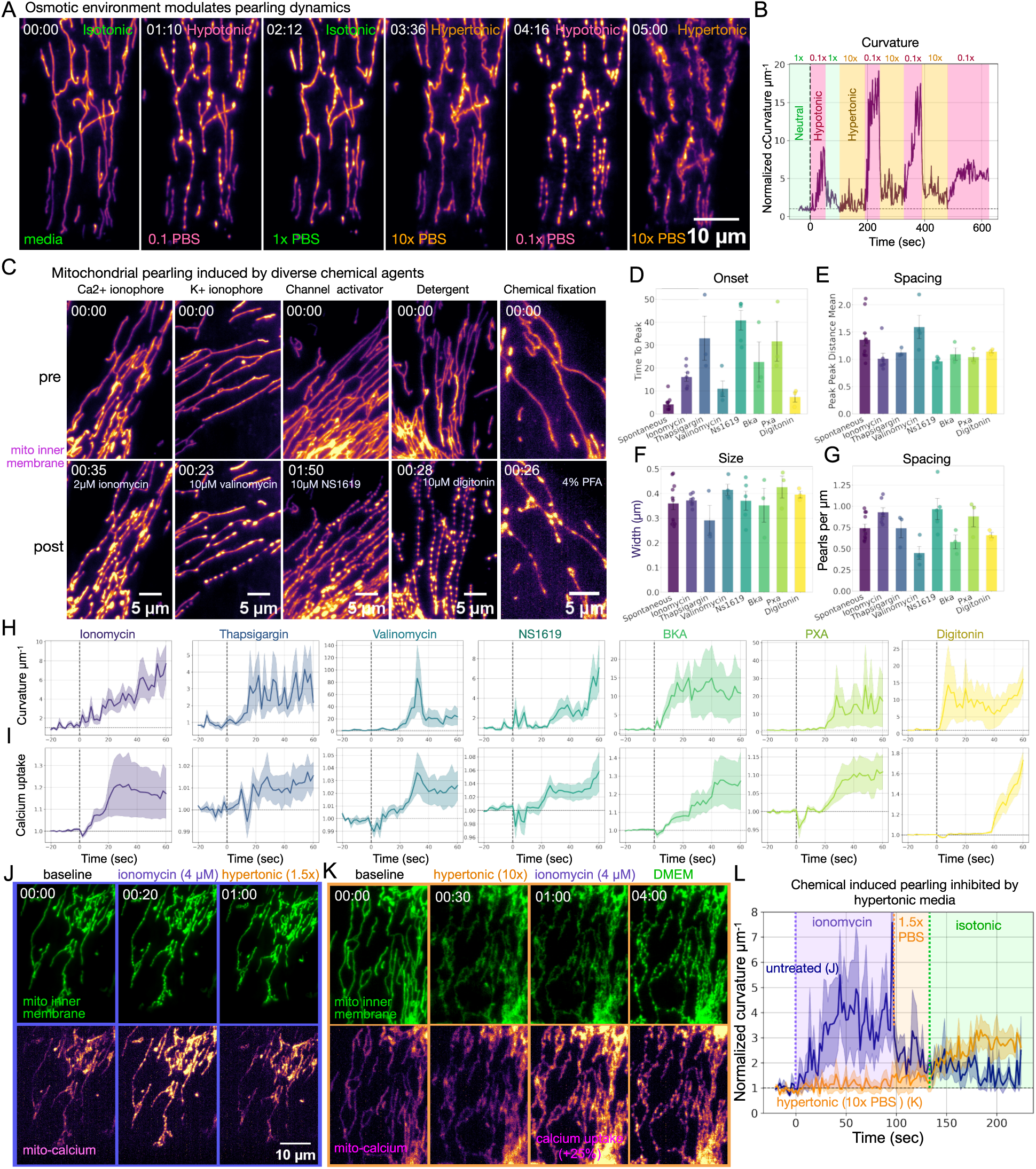
Osmotic pressure underpins chemically induced mitochondrial pearling. (**A**) Osmotic shock of primary human fibroblasts. Media is repeatedly swapped between isotonic (DMEM), hypotonic (0.1x PBS) and hypertonic (10x PBS) conditions. (**B**) Quantifying pearling based on normalized curvature per µm over time during different osmotic shock regiments corresponding to (A). (**C**) Classes of chemical inducers of mitochondria pearling encompassing calcium and potassium ionophores, potassium channel activator (BKCa), membrane permeabilization, and chemical fixation. (**D-G**) Comparison of chemical agents along with spontaneous pearling of primary human fibroblasts for the (D) time to the peak of pearling curvature (E) the distance between the centers of neighboring pearls at the peak of pearling, (F) the cross-diameter width of pearls, and (G) the number of pearls per micron of tubule length before the onset of pearling. (**H**) Panel of mitochondrial curvature per µm normalized to first time points over time. Dotted line indicates the point of injection of the chemical agent. (**J**) Rescue of tubular morphology induced by ionomycin (2 µm) by addition of 1.5x hyper-osmotic PBS to the media. (**K**) Inhibition of ionomycin-induced pearling (2 µM) by preconditioning the cell with hypertonic media (10x PBS). Three minutes after ionomycin addition, media is swapped back to untreated isotonic DMEM. (**L**) Intensity-based curvature per µm normalized to first time points over time for primary human fibroblasts without pre-treatment (blue) or placed in hypertonic 10x PBS (orange). Cells then underwent chemical-induction by 4µM ionomycin and recovery by either 1.5x hypertonic solution or isotonic media. Vertical dotted lines indicate injection of 4 µM ionomycin (purple), injection of hypertonic 1.5x PBS (orange), or media swap to isotonic media (green). n=7 cells per group.

To determine whether a specific ion or channel regulates mitochondrial osmotic pressure, we performed perturbation experiments using several chemical agents capable of altering mitochondrial ionic balance (**Table 1; SI Appendix, Fig. S4**). Calcium and potassium ionophores, channel activators, detergents, fixatives, and compounds with more complex target sets such as bonkeric acid (BKA), and phomoxanthone A (PXA) all induced cell-wide mitochondrial pearling within a minute of treatment (**Fig. 2C; Movie S16**). To confirm that these agents acted through their respective ionic species, we co-labeled cells with mitochondrial calcium marker Mito-R-Geco1. As expected, the calcium ionophore ionomycin, as well as thapsigargin, induced an uptake in mitochondrial calcium that preceded the onset of pearling. The potassium ionophore, valinomycin, and the BKCa channel activator, NS1619, caused cell-wide pearling that did not always accompany calcium uptake (**SI Appendix, Fig. S3H**), suggesting multiple ion species can induce mitochondrial pearling. Interestingly, close examination of the intermediate time points of NS1619, BKA, PXA and valinomycin immediately before the formation of pearls, revealed that mitochondria tubules entered a flaccid shape identical to hyperosmotic shock (**SI Appendix, Fig. S3G; Movie S17**). These compounds may have caused hyperosmotic pressure on mitochondria which later pearled when recovering back to isotonic balance. The membrane permeabilization agent, digitonin, resulted in the fastest onset of global pearling (**Fig. 2D; right-most graph; Movie S18**) which can be explained as an increase in osmotic pressure as water followed the ionic gradient into the cell with its plasma membrane degraded. Further, the widely used fixative paraformaldehyde (PFA, 4%), caused cell-wide pearling, consistent with earlier reports of it being osmotically active (42) (**SI Appendix, Fig. S3I; Movie S19**). Pearl formation was inhibited by fixation with a combination of 3% PFA and >0.5% glutaraldehyde (GA), but this was cell type dependent. (**SI Appendix, Fig. S3J; Movie S20**). This inhibition is likely due to the faster fixation rate of GA (see **SI Appendix, Supporting Text**). Other reported inducers of mitochondrial pearling not tested here include histamine, tunicamycin, A23187, and glutamate, which similarly activate intracellular ionic flux (19, 43–45). Pearls induced by these diverse chemical inducers had similar width, peak-to-peak distances, and pearls per µm within the initial onset of pearl formation as seen in spontaneous pearling events (**Fig. 2D-G**). Further, all of these chemical agents led to decreased tubule length (**SI Appendix, Fig. S4C**), again reminiscent of class I spontaneous events.

**Table 1:** List of chemical agents that induce cell-wide mitochondrial pearling transitions. . NA indicates the combination was not tested or is not experimentally plausible.

| Pearling inducer | Concentration ($\mu\text{M}$) | Time of onset (sec) | Known target | FCCP dependent | Hyperosmotic recovery |
| --- | --- | --- | --- | --- | --- |
| ionomycin | 2-20 | 15-45 | calcium ionophore | no | yes |
| thapsigargin | 0.2-10 | 20-50 | inhibitor of calcium ATPase (SERCA) pump | no | yes |
| valinomycin | 10 | 20-50 | potassium ionophore | no | yes |
| NS1619 | 10-30 | 60-90 | BKCa channel activator | yes | yes |
| PXA | 10 | 30-60 | fungal toxin | no | yes |
| BKA | 10 | 15-45 | ATP/ADP translocase and mPTP inhibitor | no | yes |
| digitonin | 10 | 5-35 | membrane permeabilization | no | yes |
| PFA | 4% | 30-60 | chemical fixative | NA | yes |

Confirming that all of these compounds act via osmotic pressure, adding 1.5x PBS (420 mOsm/L) into the media containing the chemical agent rescued pearled morphology (**Fig. 2J; Movie S21**). Finally, and most importantly, when cells were placed in a hyperosmotic solution of 10x PBS (2800 mOsm/L), the calcium ionophore ionomycin still caused calcium uptake in mitochondria but tubules remained flaccid, without any visible pearling or rise in membrane curvature (**Fig. 2K-L; Movie S22**). Only after removal of the hypertonic media did the mitochondria transition to pearls. This is direct evidence that osmotic pressure mediated the effect of ionic flux. This shared property amongst diverse chemical inducers of pearling indicated that it is unlikely that a single molecular ligand-binding mechanism can explain the shared dynamics of such experiments. Instead, osmotic pressure itself, being the common feature in all cases, is the driving force of these pearling events.

### Mitochondrial pearling can be induced with single-mitochondrion precision

To experimentally manipulate pearling dynamics of individual mitochondria, we exposed cells to focused laser light (0.3 mW for 200 milliseconds) pinpointed to individual mitochondrial tubules (**Fig. 3A**). This stimulation photobleached fluorophores within a <1 µm diameter (**SI Appendix, Fig. S5A**), while inducing pearled morphology in all mitochondrial tubules connected to the tubule targeted by the laser, but not in tubules unconnected to the stimulation point. In primary fibroblasts, with long and fused mitochondrial tubules, laser stimulation induced pearls as far as 140 µm away from the point of stimulation, again highlighting that connected components of mitochondrial network are biophysically coupled (**Fig. 3B; Movie S23**). Stimulating a single point in a fully fused mitochondrial network induced pearling across the entire cell (**Fig. 3C; Movie S24**). As with spontaneous pearling, laser-induced pearls recovered a cylindrical morphology on average 15 seconds post laser stimulation (**Fig. 3D**), and did on occasion revert back to a pearl morphology within 60 seconds post-stimulation (**SI Appendix, Fig. S5B; Movie S25**). Pearls induced by laser stimulation tended to have farther peak-to-peak distances than spontaneous events but otherwise maintained similar pearl structure and dynamics to spontaneous events (**Fig. 3E**). Colabeling with Mito-R-GECO1 revealed a rise in mitochondrial calcium uptake that sometimes preceded the onset of pearls (**SI Appendix, Fig. S5D; Movie S26**). These pearling events were all marked by retraction of branching tubules, suggesting a shared mechanism to class I spontaneous pearling events (**Fig. 3F**). Like chemical inducers of pearling, laser stimulation did not pearl mitochondria when cells were placed in hypertonic media (**Fig. 3D**), suggesting osmotic pressure is also the downstream driver of pearling resulting from laser-stimulation.

**Figure 3.**
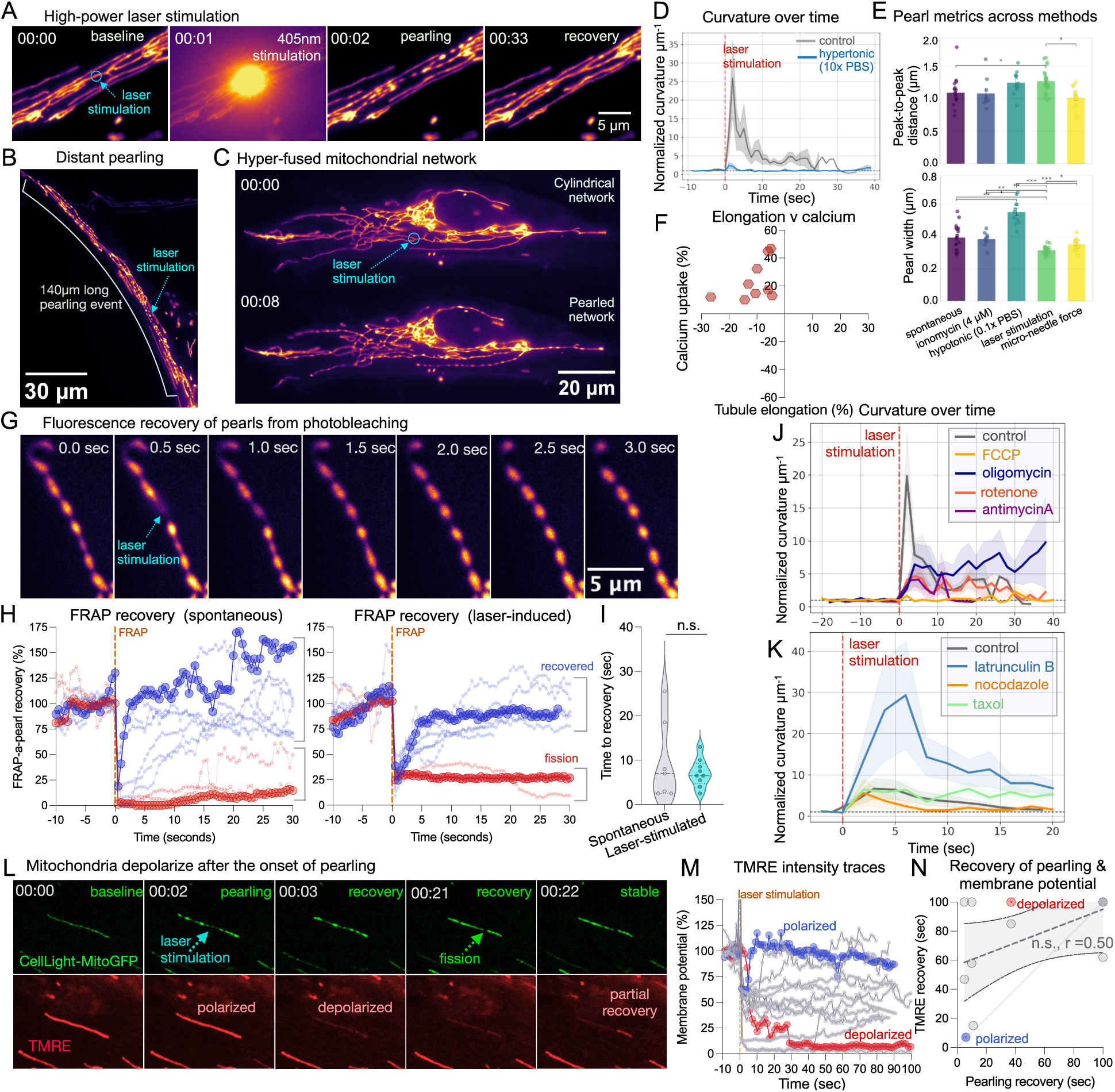
Laser stimulation induces pearling of individual mitochondrial tubules. (**A**) Pearling induced on primary fibroblast by a pinpointed 405nm stimulation laser. The 405 nm laser was fired at 20mW (0.3mW reaching sample) for 200 milliseconds pointed directly on the mitochondrial tubule (blue arrow). (**B**) Primary fibroblast with elongated mitochondria tubule pearled by laser stimulation. (**C**) Hyper-fused mitochondrial network of a primary fibroblasts before and after laser stimulation to a region marked by a blue circle. (**D**) Curvature per µm over time for laser stimulated untreated cells (n=22, gray) and cells placed in 10X PBS (n=15, blue). (**E**) Comparison of peak-to-peak distance between pearls and pearl width (minor-axis) amongst different methods to induce mitochondrial pearling. One-way anova model used. * = p < 0.05, ** = p < 0.01, *** = p < 0.001. (**F**) Scatter plot of calcium uptake vs tubule elongation amongst laser-stimulated pearling events. Red dots mark events under which shrinking occurred. n = 10 events. (**G**) Spontaneous mitochondrial pearls depleted with laser and fluorescence recovery tracked over time. (**H**) Fluorescence recovery after photobleaching of individual pearls of spontaneous and laser stimulated pearling events (n = 10 events/group). Each line indicates an individual event. Blue indicates recovery, while red indicates minimal recovery. (**I**) Comparison of FRAP recovery times between different methods of induction, Mann-Whitney non-parametric t-test. (**J**) Curvature per unit µm comparing electron transport chain inhibitors, n=21 oligomycin (1 uM) events, n=18 FCCP (4 µM) events, n=18 rotenone (5 uM) events, n=18 antimycin A (5 uM) events. Red dotted line indicates the point of laser stimulation. All data normalized to the average of the first five timepoints for each treatment group. (**K)** Curvature per µm over time for laser stimulated untreated cells, latrunculin B (1 µM), nocodazole (200 nM), and taxol (100 nM) (n=8-10 events per treatment). Data normalized to the first three timepoints of each treatment group. (**L**) Representative time series of membrane potential collapse and recovery after laser-stimulated pearling. CellLight MitoGFP labels the IMM and mitochondrial matrix. TMRE labels mitochondrial membrane potential. (**M**) Normalized TMRE intensity over time. Each line indicates an individual event. Blue indicates an event that recovered back to baseline intensity. Red indicates an event which does not recovery within the duration of imaging. (**N**) Spearman correlation between time for pearls to recovery back to cylindrical shape and time of TMRE signal to return to stable signal. Red and blue dots match individual events highlighted in (M). Faded gray line indicates an ideal correlation of 1.0. Dotted line indicates linear regression with 95% confidence lines.

### Mitochondrial matrix connection is maintained during pearling

We confirmed material exchange between the matrices of adjacent pearls by FRAP in individual pearls (**Fig. 3G**). Fluorescence recovery within photobleached spontaneous and laser-induced pearls took 9.6 seconds and 7.1 seconds, respectively (p = 0.98), (**Fig. 3H-I**). Most pearls (>70%) recovered from depletion, indicating that pearled mitochondria maintain matrix flow between connected inner membranes. The few pearls that did not recover were all located at the tips of mitochondrial tubules, had independent movements from the adjacent tubule, and thus likely underwent complete fragmentation during the pearling event.

### ETC activity and cytoskeleton structures support pearl formation and recovery

To uncover cellular properties regulating mitochondrial pearling, we applied a series of functional markers and chemical inhibitors and then induced pearling with laser stimulation (**SI Appendix, Tables S4-5**). As with other inducers of pearling, depolarizing mitochondria with FCCP reduced pearl formation after laser stimulation by 94% (**Fig. 3J; Movie S27**). Similarly, inhibiting electron transport chain (ETC) complex I, III, and V with rotenone, antimycin A, or oligomycin, mitigated pearl curvature by 76%, 79%, and 59% respectively. Note, that these ETC inhibitors independently raised baseline curvature by 2-fold before laser stimulation (i.e. caused partial pearl formation) (**SI Appendix, Fig. S5C**). Overall, depolarization with FCCP had the most dramatic reduction in formation of pearls by laser stimulation. While untreated cells took on average 10 seconds to recover back to a uniform morphology, oligomycin-treated cells delayed recovery to 35 seconds (p<0.01). This may be because inhibiting the ATP synthase causes mild hyperpolarization on the inner membrane (46), or due to off-target effects on actin polymerization (47).

Similar to oligomycin, laser stimulation of mitochondrial tubules following depolymerization of the actin cytoskeleton with latrunculin B, resulted in a permanent transition to pearl morphology that did not recover back to the cylindrical shape (**Fig. 3K; Movie S28**). These pearls progressively accumulated into larger pearls as was seen in the downstream chemically-induced pearls. In contrast, with microtubule polymerization inhibited, pearls had reduced curvature by ∼10% and recovered back to the baseline state 10 seconds faster than controls. Further, the microtubule stabilizing agent, taxol, mimicked the behavior of latrunculin B, preventing recovery to a uniform morphology. This suggested that actin was necessary for class I pearling events to re-extend back to a cylindrical morphology. Taken together, these inhibitors revealed that both electron transport chain activity as well as cytoskeleton structure influenced pearling dynamics.

### Mitochondria transiently depolarize after the onset of pearling

Colabeling with the membrane potential marker, TMRE, revealed that the IMM depolarized after the onset of pearling and recovered back to baseline after 70% of laser-induced pearling events (**Fig. 3L-M; Movie S29**). This recovery often occurred on specific portions of the original tubule that had separated during pearl formation. As previously reported, pearl shapes recovered back to a cylindrical shape faster than the recovery of membrane potential (**Fig. 3N**) (16). Interestingly, membrane potential recovery was necessary for a second induction of pearling on the same tubule by laser stimulation. Overall, laser-induced pearling matched the behavior of DRP1-KO pearls (16), in that both required ETC activity and resulted in transient depolarization of the inner membrane. For further characterization of laser-induced pearling see **SI Appendix, Supporting Text**.

### STED reveals swollen cristae ultrastructure in mitochondrial pearls

Using super-resolution STED imaging, we characterized mitochondrial cristae ultrastructure of pearls. The cristae of laser-induced pearls (class I) under STED resembled the helicoidal ramp structures to the Mic60 complex maintaining crista junctions (48, 49) (**Fig. 4A; Movie S30**). Live-cell STED imaging confirmed a 20.4% increase in cross-sectional area and 70.7% increase in inter-cristae distance at the onset of laser-induced pearling (**Fig. 4B-C**). Pearls were connected by a continuous inner membrane and often contained nucleoids (**Fig. 4D-E**). However, pearls were also seen that did not contain nucleoids.

**Figure 4.**
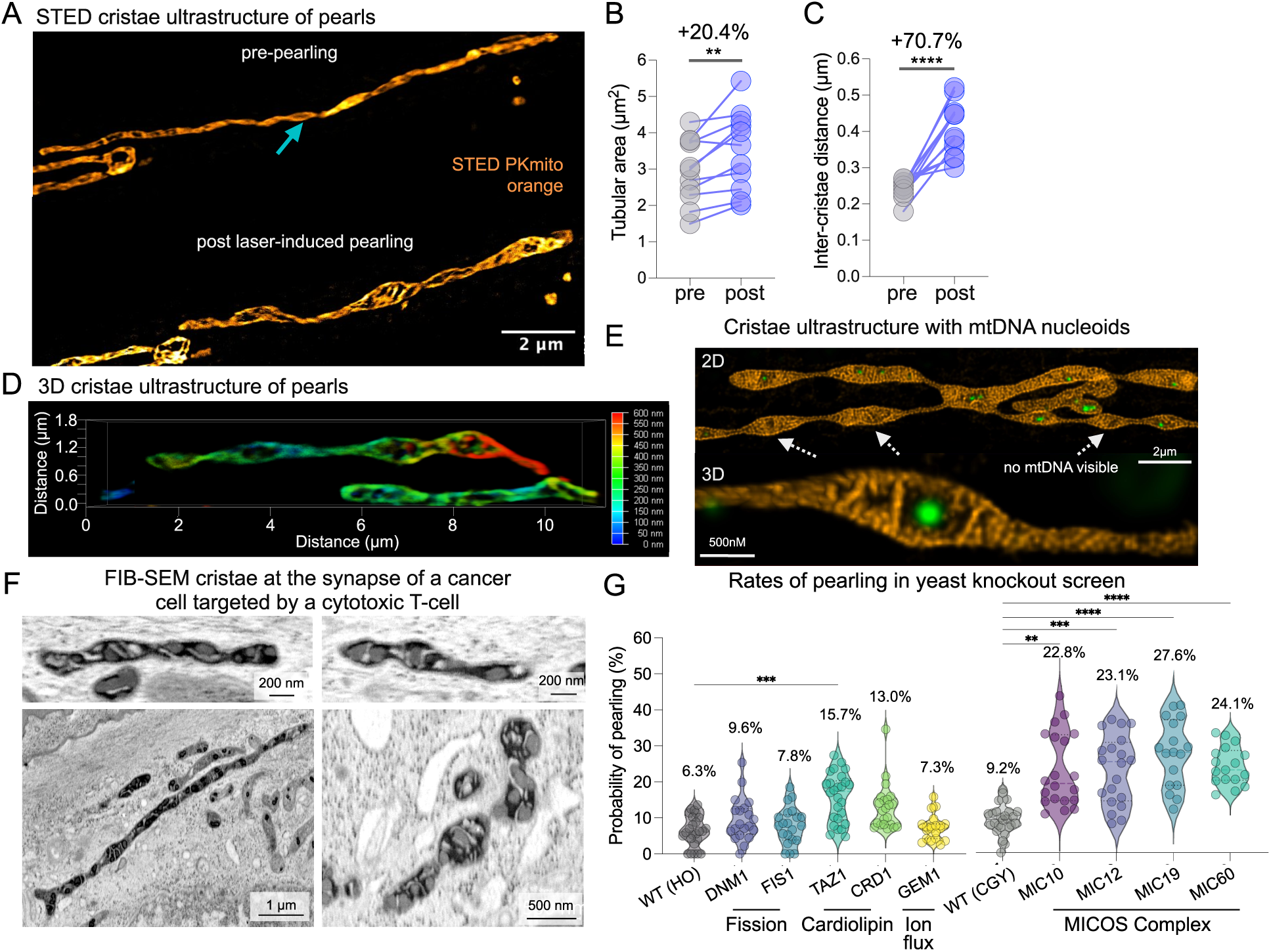
Mitochondrial cristae ultrastructure during pearling. (**A**) Live-cell stimulated emission depletion (STED) microscopy of mitochondrial cristae labeled with PKmito Orange dye, before and after laser stimulation of mitochondrial tubule. Two-way Anova, ** = p < 0.01. (**B**) Comparison of tubular area before and after laser-induced pearling. n=11 mitochondria. (**C**) Inter-cristae distance before and after laser-induced pearling. n=11 mitochondria. Two-way Anova, **** = p < 0.0001. (**D**) Cristae architecture in 3-dimensions using tauSTED. Depth-encoded color mapping. (**E**) Deconvolved STED imaging of mitochondrial inner membrane (orange, PKmito Orange) along with mitochondrial DNA (green, Sybr Gold) during laser-induced pearling event. An example 2D image is shown in the top panel and a separate 3D image in the bottom panel. (**F**) FIB-SEM mitochondrial ultrastructure responding to the release perforin and granzyme from lytic granules at the synapse of an ID8 ovarian cancer cell attacked by OT-I mouse cytotoxic T-cell (see SI Appendix, Fig. S7). (**G**) Probability of a pearling occurring in a yeast cell over 100 volumes of imaging for wild-type (CGY14.31 & CGY49.77, and genetic knockouts of DNM1 & FIS1 (fission machinery), TAZ1 & CRD1 (cardiolipin synthesis), GEM1 (ionic flux), and MIC10, MIC12, MIC19, and MIC60 (MICOS complex). n=17-25 videos per group which accounts for ∼750 cells per group. Non-parametric Kruskal-Wallis test performed. ** = p < 0.01, *** = p < 0.001, **** = p < 0.0001.

Using a previously published whole-cell FIB-SEM 8-nm isotropic dataset of ID8 ovarian cancer cells attacked by cytotoxic CD8 T-cell, we identified high-electron dense (i.e. ‘condensed’ (50)) cristae ultrastructure amongst pearled mitochondria proximal to the immune synapse (**Fig. 4F; SI Appendix, Fig. S7A-C**). This particular immune context is marked by calcium waves emanating from the synaptic cleft between the T-cell and target ID8 cell, thus generating the necessary osmotic pressure needed for class I pearling events. We note that pearl shaped mitochondria were also present further away from the synapse without high-electron dense cristae (**SI Appendix, Fig. S7C**). This high-resolution data confirmed that both outer and inner membranes transition to constricted pearl shapes and both membranes remain connected across the pearled tubule.

### Mitochondrial membrane scaffolding molecules regulates pearling dynamics

The mechanical properties of mitochondrial tubules depend on membrane composition (**SI Appendix, Fig. S8A**), including cholesterol, cardiolipin, and the MICOS complex. Depleting cholesterol with atorvastatin (24hr of 50µM) blocked pearling in response to laser-stimulation (**SI Appendix, Fig. S8B-C**). Without sufficient cholesterol the mitochondrial membrane may have been too fluid to hold tension and undergo a pearling transition. On the other hand, knockout of the MICOS complex genes, MIC10, MIC12, MIC19, and MIC60 all increased the frequency of spontaneous pearling events by 13.6% to 18.4% (p<0.0001) (**Fig. 4E**). These genes support a variety of MICOS properties including the junction’s membrane curvature (MIC10), complex subunit bridging (MIC19), and inner-to-outer membrane connections (MIC60), indicating that the integrity of the entire complex stabilizes the tubular shape and limits pearling transitions. We suspect that disruption of cristae lamella folding decreases the membrane’s exceptionally high bending modulus, and thereby basal levels of tension more easily exceed its elasticity and cross the pearling instability. Removal of the cardiolipin synthesis gene, TAZ1, or CRD1, increased the frequency of spontaneous pearling events by +9.4% (p<0.001) and +6.7% (p=0.12) compared to wildtype, respectively. This can be explained by cardiolipin’s unique lipid properties in the IMM, supporting cristae invaginations in the inner membrane (51).

### Alternative mechanisms fail to explain spontaneous pearling events

Other potential mechanisms of spontaneous mitochondrial pearling include dynamin-like fission machinery, ER constriction events, ROS oxidative stress, actin waves, and mtDNA nucleoid colocalization. Several studies reported increased pearling when DRP1 and related fission machinery was removed (5, 16, 17), showing that DRP1 mediated fission is not required. In our time lapse data of budding yeast mitochondria, knockout of fission machinery DNM1 and FIS1 did not significantly increase the rate of pearling with a +3.3% (n.s.) and +1.5% (n.s.) compared to wildtype, respectively (**Fig. 4G**).

Colabeling ER and mitochondrial networks showed that ER network structures often did not form ring shapes around pearl constriction points (**Fig. 5A-C; Movies 31-33**). In spontaneous and laser-induced pearling, mitochondrial pearl constriction points formed without any nearby ER signal labeled with KDEL and calreticulin probes in 33% and 22% of pearls, respectively (**Fig. 5D**).

**Figure 5.**
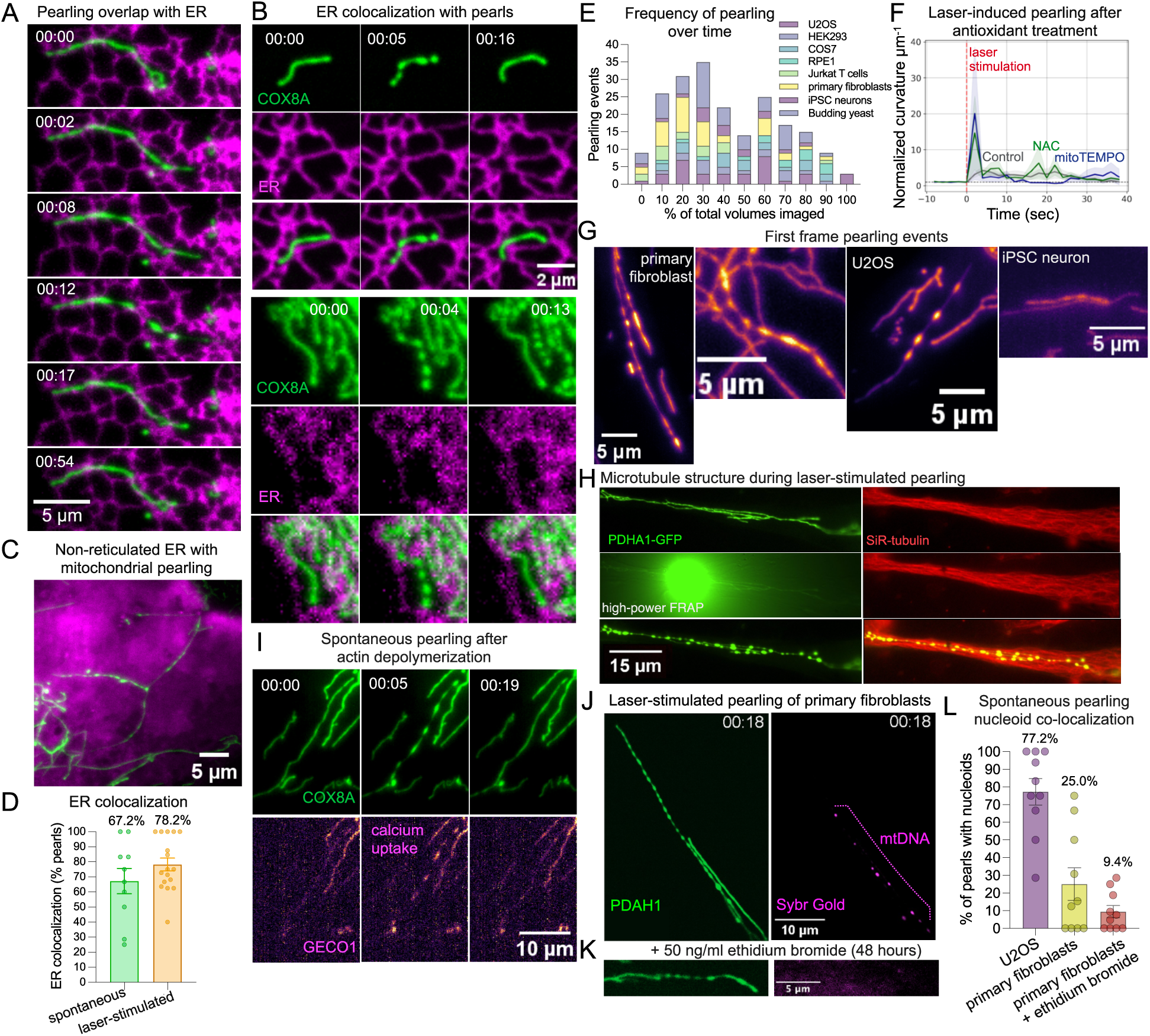
ER constrictions, oxidative stress, microtubule organization, actin polymerization, and nucleoid localization fail to regulate mitochondrial pearling dynamics. (**A-B**) Spontaneous pearling events with both mitochondria (COX8A-mEmerald) and ER (KDEL, CellLight-ER-RFP) labeled in U2OS cells. (**C**) A cloudy ER network surrounding mitochondrial pearls in a U2OS cell. Mitochondria (COX8A-mEmerald) and ER (KDEL, CellLight-ER-RFP) labeled. (**D**) Percentage of pearl constriction points with or without overlapping ER fluorescence signal from both U2OS and primary fibroblasts either from spontaneous events (green, n=10 events) or laser-induction (orange, n=17 events). (**E**) Frequency distribution of the frame at which a pearling event was first observed normalized to the total number of frames (i.e. 3D volumes) taken in a video. (**F**) Mean curvature per µm over time for primary fibroblasts treated with antioxidants 5 µM NAC and 20 nM mitoTEMPO (n=10-14 events per treatment group). Red line indicates point of laser stimulation. (**G**) Mitochondrial pearls imaged in the first frame of a timelapse video. (**H**) Laser-induced pearling with both the mitochondrial inner membrane (green, CellLight MitoGFP) and microtubules (red, Sir-tubulin) labeled. (**I**) A spontaneous pearling event observed in U2OS cells treated with 1 µM latrunculin B for 1 hr. Top panel contains the mitochondrial inner membrane (COX8A-mStayGold) and the bottom panel contains mitochondrial calcium (Mito-R-GECO1). (**J**) Mitochondrial pearls overlapping with mtDNA (Sybr Gold) induced by laser-stimulation of a primary fibroblast mitochondria in (I) untreated, and (**K**) DNA intercalating ethidium bromide (50 ng/mL) for 48 hours. IMM = CellLight MitoGFP, mtDNA = Sybr Gold. (**L**) Quantification of the number of pearls with overlapping mtDNA signal for U2OS cells (purple), primary fibroblasts (yellow) and primary fibroblasts treated with 50 ng/ml ethidium bromide for 48 hours. n=10 events per group.

To rule out the possibility of phototoxic initiation of spontaneous pearling events we examined the distribution of pearling events across all cell types (**Fig. 5E**). Spontaneous pearling did not increase with increased duration of imaging, and additionally we observed 9 events at the first volume of imaging showing that pearls were already formed in the absence of illumination (**Fig. 5G**). Further evidence that spontaneous mitochondrial pearling events are independent of phototoxic illumination is that they still occur under low-intensity label-free phase-contrast imaging (52).

Regarding the idea that ROS-related oxidative stress drives pearling, treating cells with antioxidants mitoTEMPO (20 nM) and N-acetyl cysteine (NAC, 5 µM) failed to inhibit the induction of pearling by laser stimulation (**Fig. 5F**). Additionally, mitochondrial-specific ROS markers, MitoSOX green and MitoSOX red did not increase in fluorescent intensity at the onset of laser-stimulated pearling, with a change in +0.4% and -1.2%, respectively (**SI Appendix, Fig. S5E**). Mitochondrial intermembrane ROS sensor, Hyper7 did increase by 18.3% directly after laser-stimulation but this increase occurred across the all mitochondria of the cell, and was not specific to tubules that underwent pearling (**SI Appendix, Fig. S5F-H**).

Colabeling with a microtubule marker, Sir-Tubulin, revealed that microtubules remained intact during laser induced pearling (**Fig. 5H**). Microtubule depolymerization with 10 µM nocodazole did not stop the induction of pearling by laser-stimulation (**Fig. 3K**) or by chemical-induction with 4 µM ionomycin (**SI Appendix, Fig. S5J)**. Further, actin depolymerization with 1 µM latrunculin, did not stop spontaneous pearling events (**Fig. 5I**) or pearling induced by 4 µM ionomycin (**SI Appendix, Fig. S5K**). Instead, the lack of actin inhibited pearls from recovering back to the tubule form (**Fig. 3K**).

Another explanation of pearls could be reorganization of mtDNA nucleoids to discrete spheres that create membrane bulges. In U2OS cells, 77% of pearls colocalized with mtDNA nucleoids, while in primary fibroblasts only 20% of pearls colocalized with nucleoids (**Fig. 5J-L**). Depletion of mtDNA with ethidium bromide did not stop induction of pearling by laser-stimulation (**Fig. 5K-L**).

### Micro-needle manipulation induces mechanical force-based pearling on individual mitochondria tubules

The chemical and laser induction of pearling described above can all be explained by the influx of ions resulting in mitochondrial shrinkage, swelling, and osmotic pressure. An alternative mechanism for pearling via the Rayleigh-Plateau instability could involve mechanical forces pulling on the membrane, thus explaining class II stretching-associated spontaneous pearling events. To test this idea, we applied mechanical forces to individual mitochondrial tubules using a fluorescently-labeled glass microneedle with a triple axis (X,Y,Z) 56 nm step-motor (**Fig. 6A**)(53). Bringing the micro-needle within a micron of the mitochondrial tubule induced rapid and transient pearling of the selected mitochondrion (**Fig. 6B; Movie S34**). Like laser-induced pearling, depolarizing the mitochondria with FCCP inhibited the ability of needle force to convert tubules to pearls (**Fig. 6E; Movie S35**). Attempting to further manipulate the tubule by pulling the depolarized tubule largely resulted in cutting (i.e. mechanical fission) at the point of needle manipulation. Regardless of whether the mitochondria were polarized or not, insertion of the needle into the cell often induced a rise in cell-wide mitochondrial calcium uptake, quickly followed by mitochondrial pearling across the entire cell (**Fig. 6C-D; Movie S36**). These global pearling events may be a result of actin waves caused by the introduction of the needle in the cell. Previous work has shown the calcium ionophore, ionomycin, induces actin waves followed by calcium uptake into the mitochondria and an increase the rate of pearling events (19).

**Figure 6.**
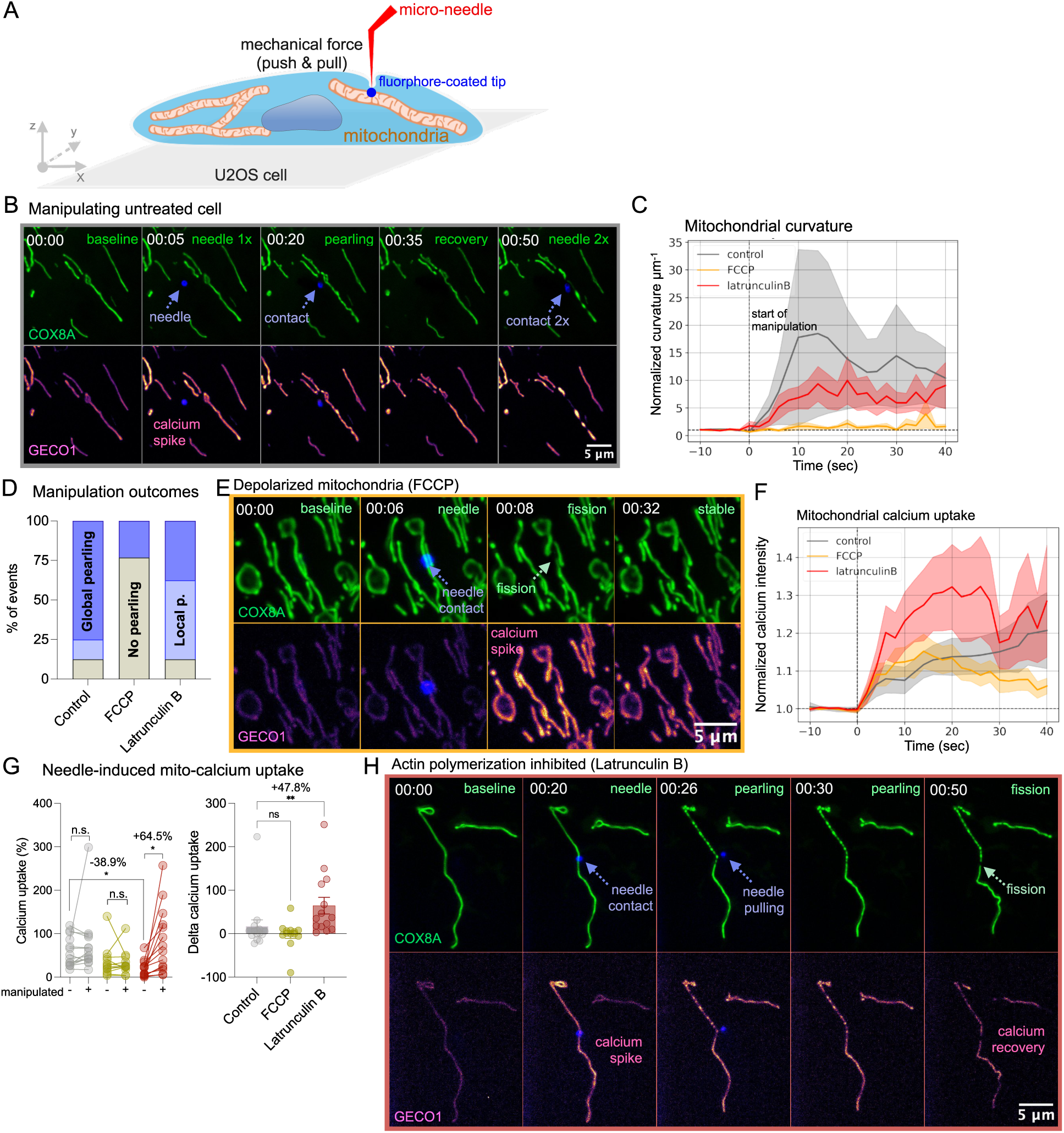
Micro-needle mechanical force triggers localized mitochondrial pearling. (**A**) Schematic of micro-needle experimental design. Red needle is manipulated in x, y, z dimensions with a 56 nm step-size control. The tip is coated with fluorescent pigment (far-red channel, labeled dark-blue in Fig.). Pico-nano newton forces are then applied to individual mitochondrial tubules with push and pull forces. (**B**) Representative time series of untreated U2OS cell mitochondria manipulated with micro-needle. Mitochondrial inner membrane labeled with COX8A-mStayGold (green). Mitochondrial calcium flux labeled with Mito-R-GECO1 (red). Needle was pushed against tubule before being removed from the cell and then reapplied for a repeated manipulation of the same tubule. (**C**) Curvature over time for single mitochondria tubules manipulated with micro-needle in untreated controls (n=14 cells), FCCP (10µM, 10-60 minutes, n=13 cells) and Latrunculin B (10µM, 30-60 minutes, n=16 cells). Thick lines represent mean weighted curvature per µm normalized to the first 3 timepoints before manipulation. Error bars indicate standard error of the mean. Dotted vertical line indicates the beginning of needle manipulation on the tubule. Curvature values are weighed per µm of distance and normalized to the mean of the first three pre-pearl timepoints. (**D**) Categorical outcome from attempted micro-needle manipulation of mitochondria. No pearling (gray) indicates that curvature of all mitochondria tubules remained stable throughout the experiment. Local pearling (light-blue) indicates pearling was uniquely induced on the tubule targeted by the needle. Global pearling (dark-blue) indicates that all other tubules in the cell underwent pearling regardless of wether it was targeted by the micro-needle. 13-16 cells per treatment group. (**E**) Representative time series of U2OS cell mitochondria depolarized with 10µM FCCP for 30 minutes and manipulated with micro-needle. (**G**) Calcium uptake derived from Mito-R-GECO1 maximum intensity values after the needle is present in the cell for non-manipulated (-) and manipulated (+) mitochondria. Right-panel shows delta calcium uptake between the mitochondria manipulated by the needle and those untouched. (**H**) Representative time series of manipulated mitochondria from a U2OS cell with actin polymerization inhibited by 10µM Latrunculin B for 45 minutes.

To overcome the needle’s tendency to cause global pearling, the actin polymerization inhibitor, Latrunculin B, was applied (1 µM for 30-60 minutes). In these cells, manipulated mitochondria tubules were more likely to undergo localized pearling specifically at the targeted mitochondria, while non-manipulated mitochondria remained stably cylindrical (**Fig. 6C-D**) and showed minimal calcium uptake (**Fig. 6G**). This further dispels the idea that actin is itself the constrictive actor that forms mitochondrial pearls. With actin inhibited, pulling with the needle caused pearling specifically on the portion of the tubule undergoing deformation (**Fig. 6H; Movie S37**). Further, in some cases force-induced pearling preceded a rise in calcium uptake by the mitochondria (**SI Appendix, Fig. S9; Movie S38**), showing that calcium on its own was not sufficient to induce pearling but required an added mechanical force on the membrane. Thus, both the internal osmotic pressure of ionic flux as well as external mechanical tension of the membrane can result in a pearling instability across the mitochondrial tubule.

## Discussion

### Mechanism of pearling induction

Our biophysical model of mitochondrial pearling describes the organelle as a network of membranes under tension that exists near a critical threshold of instability. Under sufficient tension these tubules collapse and reorganize into thinly connected spherical units. This instability is regularly crossed due to three types of physical changes: osmotic pressure resulting from ionic flux of cations into the mitochondrial matrix; changes to elasticity of the membrane such as the removal of cristae junctions; and external forces pulling on the membrane. Ultimately, all three physical changes can result in the membrane tension surpassing its bending energy (**Fig. 7**). This biophysical mechanism unifies mitochondrial pearling events observed in diverse chemical, physical, and genetic, and physiological scenarios (**SI Appendix, Fig. S10**). In physiological scenarios such as neuronal action potentials and T cell synapse formation, pearling is likely a result of ionic flux, which results in osmotic pressure on the inner membrane.

**Figure 7.**
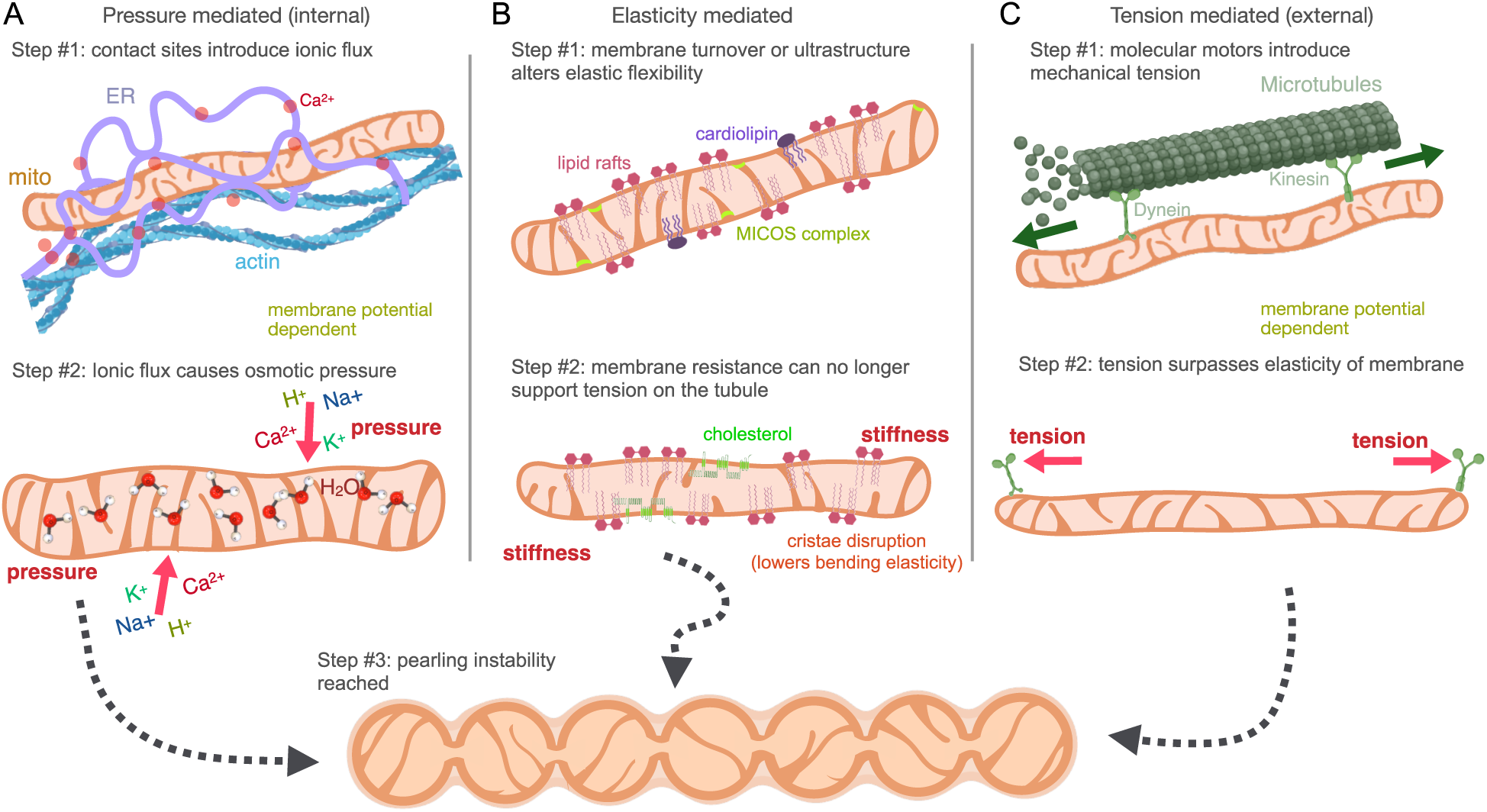
Biophysical mechanism of mitochondrial pearling dynamics. (**A-C**) Pressure, tension, and elasticity are the three main physical properties that influence the pearling instability of the mitochondria membrane. (**A**) Contact sites with other organelle systems introduce ionic flux on the mitochondrial membrane. Ionic flow into the mitochondria causes an osmolarity difference between the inside of the mitochondria causing an influx in water. Osmotic pressure increases tension on the membrane which exceeds its bending elasticity and collapses the mitochondrial tubule into pearls. (**B**) Alteration to components of the mitochondrial membrane such as cholesterol, cardiolipin, and cristae organization change the elasticity of the membrane and can collapse the tubular mitochondria into pearls. (**C**) Molecular motors traffic mitochondria and introduce external mechanical tension through pulling forces on the membrane. Mechanical tension exceeds the bending elasticity of the membrane and collapses the mitochondria tubule into pearls, following a Rayleigh-plateau ‘pearling’ instability.

### Spontaneous pearling triggered by multiple physical forces

With this biophysical framework, the two different types of spontaneous pearling we have uncovered, stretching and calcium associated pearling, respectively, can be driven by two distinct biophysical causes. Stretching can be produced by cytoskeletal motor proteins tethered to mitochondria and pulling the membrane (54). This is directly validated by activation of these motors with optogenetically-activated motors, which results in force-dependent elongation of mitochondrial tubules followed by network wide mitochondrial pearling events within the stimulated cell (55). On the other hand, spontaneous events that coincide with calcium uptake and membrane shrinkage may be explained by calcium waves propagating through the cell that are soaked up by mitochondria’s buffering capacity (56) increasing internal osmotic pressure. Our results also indicate that changes to mitochondrial membrane components, particularly the MICOS complex proteins regulating cristae junctions, altered the mitochondria’s capacity for pearling. Disruption of cristae ultrastructure reduces the membrane’s exceptionally high bending modulus (21) thereby increasing the frequency of spontaneous pearling. Only by characterizing the state of individual mitochondrial tubules in terms of osmotic pressure, mechanical tension, and membrane elasticity can one predict a pearling transition.

### Electrochemical coupling supports mitochondrial pearling

Our findings indicate that the membrane polarization maintained by the IMM is a primary regulator of pearling transitions. This suggests that the flux of ions and water that build tension on the IMM are supported by the electrical coupling of the IMM. Future work will be required to determine how the inner membrane’s electrochemical coupling is capable of regulating pearling events. We postulate that membrane polarization may induce a conformation of lipids that supports increased tension on the membrane. Without the electrochemical gradient, mitochondrial membranes become “soft” with disorganized lipids and thereby unable to maintain higher tension states. Further, like a neuronal action potential, the electrical pulse resulting from membrane depolarization should be accompanied by a mechanical wave that propagates across the membrane (57). This electrically-driven mechanical wave may drive the structural reorganization necessary for pearl formation. Alternatively, mitochondrial membrane polarization may be necessary to support ionic flux between mitochondria and the cytoplasm. Without an internal ionic gradient, depolarized mitochondria will not undergo class I pearling events. However, direct osmotic shock or ionophores still result in class I pearling events despite a lack of mitochondrial polarization by driving water influx via extracellular ion gradients.

### Functional utility of mitochondrial pearling

Beyond generating ATP, one core function of mitochondria is buffering ions within cells. Mitochondrial membranes contain many ionic pumps regulating the flux of H+ (ATP synthase), Ca2+ (calcium uniporter, MCU), K+ (mitoKATP, mitoBKCa, mitoKv, mitoTASK, mitoIKCa), Na+ (mitoSLO2), Mg2+ (MRS2), and Cl- (CLIC4&5, Ano-1). Given that the presence of each channel varies from tissue to tissue, but we observe pearling in diverse cell types across evolutionary distinct species, hints that it is unlikely for one specific ion channel to drive all pearling events. However, it is still possible that many of these pores evolved to mitigate or potentiate pearling events for cellular homeostasis.

We have yet to establish how pearling affects mitochondrial function, particularly energy production. Current readouts of bioenergetic output, namely Seahorse extracellular flux analyzer, measures oxygen consumption on the order of minutes and pearling occurs on the order of seconds. Future instrumentation may enable access to energy production on the same timescale as a pearling event. Additionally, *ex vivo* measurement of mitochondrial pearling is limited by the lack of experimental methods that extract mitochondria from the cell without conversion of mitochondrial tubules into fragmented pearls. We hypothesize that energy production will follow a biphasic effect with increased OXPHOS as the IMM increases in tension, followed by a precipitous decline in OXPHOS upon crossing the pearling instability. Supporting evidence is the fact that membrane potential collapses after the onset of pearling (**Fig. 3H-I**). This leads us to believe that in energetically demanding cells, such as muscle cells, mitochondria are driven to the threshold of membrane tension to maximize energy production. Pearling, in this context, represents a physical barrier or breakpoint limiting energetic output. In other cell types such as neurons or cardiomyocytes, mitochondria’s primary function is to buffer ions to help potentiate action potentials periodically moving across the cell (56, 58). In this context, pearling dynamics may be a necessary event to support ionic buffering with ionic channels tuned to ensure rapid pearl recovery for periodic neuronal activity.

### Connection between mitochondrial pearling to fission

We hypothesize that Drp1 knock-out cells have an increased rate of pearling (5, 16) because in its hyperfused network, pearling events will spread through the whole network, instead of the typical event occurring along a relatively isolated segment of mitochondria. Moreover, without fission, hyperfused mitochondria may enter a higher tension regime, similar to senescent fibroblasts, and thereby more readily cross a pearling instability. Molecular fission machinery may thereby serve to mitigate the physical coupling of pearling, isolating these transitions to smaller regions of the cell.

## Conclusion

Our experimental results indicate that mitochondrial pearling is governed by the biophysical forces of elasticity, pressure, and tension. This mechanism bridges mitochondria dynamics with ionic flux, molecular motors, and cellular energetics. Pearling is thereby a prime example justifying mitochondria’s role as the central processor of the cell (59) simultaneously integrating both biochemical and biophysical signals. Other mitochondrial dynamics such as fission and fusion were previously implicated in aging and age-related disease (60). Mitochondrial pearling may similarly advance our understanding of key health transitions. With that in mind, this paper establishes a unified biophysical model to understand the driving forces of mitochondrial pearling.

## Data and code availability

The code and processed data associated with this manuscript is available at https://github.com/gav-sturm/Mitochondrial_Pearling. Further access to raw data files is available upon reasonable request to the corresponding author.

## Author Contributions

G.S. conceptualized the study, designed and ran the experiments, analyzed the data, and wrote the initial draft of the manuscript. K.H. assisted in project conceptualization and experimental design. A.Y.T.L developed image analysis software and assisted in analysis. C.R. and S.D. developed and assisted in microneedle experiments. D.I. assisted in running laser stimulation experiments. M.C. generated iPSC neurons. K.M.T. assisted in experimental design.

W.L. designed and assisted in T cell activation experiments. B.R. and A.W. generated yeast knockout strains. A.R. generated and processed FIB-SEM Data. J.C.L assisted in project conceptualization and protocol development. W.M. and S.M. secured funding and assisted in project conceptualization. All authors reviewed and edited the manuscript.

## Acknowledgements

This work was funded by NSF GRFP, HFSP grant RGP0038/2021 (SM and WFM), NIH grant R35 GM130327 (WFM), the Center for Cellular Construction supported by NSF grant DBI1548297 (SM and WFM). We further thank the Calico Life Sciences internship program for the initiation of this project.

## Supplemental Text

### Chemical fixation induces mitochondrial pearling artifacts

Paraformaldehyde is osmotically active (1). Others have found that addition of glutaraldehyde can mitigate this artifact through its faster fixation rate (2–4). It is worth noting that in addition to a pearling artifact, the reverse ‘hyper-osmotic’ artifact of flaccid tubules of mitochondria is also visible at higher levels of glutaraldehyde (**SI Appendix, Fig. S3D-E**). To maintain ideal ultrastructure it is necessary to perform a careful titration of PFA:GA or fixing the sample with high-pressure freezing or cryo-fixation (5). With this in mind, earlier reports of mitochondrial pearling in tissue samples (6, 7) may have been artifacts of the osmotic pressure exerted by 4% PFA during perfusion and that specific experimental groups may contain mitochondria more susceptible to osmotic pressure.

### Endogenous fluorescence predicts pearling response to laser stimulation

Dim-emission detection of the 488-525 nm channel revealed an endogenous fluorescent signal coming from mitochondrial inner matrix that accumulated after the onset of both spontaneous and laser-stimulated pearling events (**SI Appendix, Fig. S7A-B**). This endogenous molecule that fluoresces in the green channel accumulates in the mitochondrial matrix after pearling occurs and sometimes recovers. This autofluorescent signal was amplified by depolarization with FCCP (4 µM) or cholesterol depletion by atorvastatin (50 µM) (**SI Appendix, Fig. S7C**). The identity of the endogenous fluorophore could be the endogenous metabolite FAD, whose autofluorescent properties have been widely reported (8–10). Whatever its identity, this fluorescence predicted whether an individual mitochondrial tubule would pearl from laser stimulation (**SI Appendix, Fig. S7D**).

## Supplemental Methods

### Mammalian cell culture

Cell lines include U2O2, HEK293, COS7, RPE1, human primary fibroblasts (hFB12, hFB13), Jurkat cells, Raji cell, primary human CD8 T cells, and iPSC-derived neurons (**Table S1**). All cells were maintained in normal 37°C incubator conditions. Immortalized cells were grown in DMEM (GlutaMax, 25mM glucose) supplemented with 10% FBS, passaged weekly, and maintained for less than 20 passages. Primary human fibroblasts were grown in DMEM (GlutaMax, 5.5mM glucose). Replicative senescent fibroblasts were generated by serial passaging of primary fibroblasts until growth arrest, which took 80 days fo growth over 50 passages and population doublings.(11)

Non-adherent Jurkat cells were grown in RPMI media supplemented with 10% FBS. Cells were transduced with lentivirus labeled with COX8A-mitoStayGold using a spin-transduction protocol for 1.5 hours at 2400 rpm. Plating was performed in an 8-well chamber slide coated with 10 µg/ml fibronectin and 5 µg/ml ICAM-1 for 3 hours at 37°C. Jurkat cells were then co-cultured with Raji cells pulsed with the Staphylococcal enterotoxin superantigen (SEE-pulsed)(12) and labeled with Sybr Gold (1:10,000x) in the chamber for 30 min at 37°C before imaging.

Primary human CD8 T cells were grown in X-VIVO 15 media (Lonza #04-418Q) with gentamicin and phenol red, supplemented with 5% FBS, 4 mM n-cysteine, 55 mM 2-mercaptoethanol (Thermo #21985023). Primary cells were activated by three day incubation with DynaBeads Human T-Activator CD3/CD28 (Thermo #11131D). Beads were magnetically removed and cells imaged with 8 hours of activation. For synapse formation, Raji cells labeled with Sybr Gold (1:10,000x for 30 min) were co-cultured with primary T cells labeled with PKmito Orange (500 µM for 1hr). Synapse formation between T cells and Raji cells were found by scanning imaging plates for adjacent cells with distinct fluorescent labels.

iPSCs were cultured in mTESR Plus media (Stem Cell Technologies #100-0276) on hESC qualified matrigel (Corning #354277) coated tissue culture dishes. Differentiation to ventral midbrain identity was adapted from a previously described method (13) with modifications for 3D embryoid body based differentiation instead of 2D differentiation. On Day 0, iPSCs were dissociated into single cells with accutase and seeded into ultra low attachment 96 well plates at 5K cells per well to form embryoid bodies in Neural Differentiation Media (1:1 Neurobasal (Thermo #A3582901)/Advanced DMEM F12 (Thermo #12634010), N2 (Thermo #17502048), B27 -vitA (Thermo #12587-010), Glutamax (Thermo #35050-061), Sodium Pyruvate (Thermo #11360070),

NEAA (Thermo #11140050), Pen/Strep (Thermo #15140-122) and 55 µM b-ME (Thermo #21985-023) supplemented with 10uM SB431542 (Selleck Chemicals #S1067), 100 nM LDN193189 (Selleck Chemicals #S2618), 0.75 µM CHIR99021 (Thermo #S1263), 500nM SAG (Thermo #S6384), 100 µM Ascorbic Acid (Sigma #A4034-100G) and 10 µM Y-27632 (BioTechne #1254).

After 2 days media was exchanged to remove Y-27632. On day four of differentiation media was exchanged and CHIR99021 concentration was increased to 7.5 µM. On Day 7 of differentiation embryoid bodies were collected and transferred to 10 cm low attachment plates using wide bore pipet tips and incubated for the remainder of differentiation on an orbital shaker at 40 rpm. Media was changed to Neural Differentiation Media supplemented with 7.5 µM CHIR99021, 500 nM SAG, and 100 µM AA until Day 10. On Day10 CHIR99021 and SAG were removed and 0.5 ng/mL TGF-b (Bio-Techne #243-B3-002/CF) and 200 µM dbcAMP (Sigma #D0627-1G) was added. On Day 12 media was exchanged and supplemented with 10 µM DAPT (BioTechne #2634/50). On Day 15 media was exchanged to the same conditions with an additional supplement of 10 ng/ml BDNF (Bio-Techne 11166-BD-01M) and GDNF (Bio-Techne #212-GD-01M/CF). Media was maintained the same changing every 3-4 days until dissociation. On Day 20-22 of differentiation EBs were collected and dissociated at 37°C in Accumax(∼15 EBs/ml of Accumax). EBs were incubated at 37°C with agitation. Following 15 mins EBs were pipetted with p1000 narrow bore tips to check state of dissociation. If still intact then EBs were returned to the shaker. Following 30 mins of enzymatic digestion 25 µg/ml DNase1 was added and EBs were manually triturated 5-10 times to begin manual dissociation. If EBs begin to dissociate then they are manually triturated ∼10-20 times, if not they are returned to the shaker for an additional 10 mins. EBs were dissociated by manual trituration into single cells with one volume of media. After dissociation neurons were pelleted by centrifugation and then filtered through a 30 µm cell strainer before plating to remove clumps. Neurons were plated in Neuronal Maintenance Media (Neurobasal Plus, B27 plus, Glutamax, Pen/Strep and 55 µM b-ME) supplemented with 10 ng/ml BDNF, 10 ng/ml GDNF, 100 µM Ascorbic Acid, 200 µM dbcAMP, 10 µM DAPT and 20 mM each of Uridine (Sigma #U3750) and Fluorodeoxyuridine (Sigma #F0503) for 1.5 weeks. Neurons were plated on 8-well chamber slides pre-coated with poly-ornithine, laminin and fibronectin. For imaging experiments, neuronal cultures were labeled with PKmito Orange (500 µM for 1 hr) and CalBryte^TM^(1 µM for 1hr) followed by a 3x media wash and a final addition of probenecid (1 mM).

### Yeast culture

Budding yeast were grown in synthetic complete medium supplemented with 2% glucose and 2% methyl-alpha-D-mannopyranoside (MMP, Sigma #M6882) to the stationary phase. Cells were plated in 8-well glass-bottom chamber slides (CellVis #C8-1.5H-N) pre-coated overnight with 1:100 concanavalin A. Yeast knockout strains are listed in **Table S2**.

### Fluorescent Labeling

All fluorescent labels used in the current study are listed in **Table S3**. Transiently transfection of cells for Mito-R-GECO1 plasmid (Addgene #46021) was performed overnight using 0.2 µl lipofectamineTM 3000 (Thermo #L3000008) with 250 ng of plasmid DNA and 0.2 µl P3000™ reagent following the manufacturer’s protocol. Stable cell line of COX8a-mEmerald was made by transfection with 1 µg of DNA using the SE Cell Line Nucleofector Kit (Lonza #V4XC-1032) following the manufacturer’s protocols. Transfected cells were cultured on collagen-coated plates to facilitate the recovery process. After two days, cells were selected with 1mg/ml Geneticin Selective Antibiotic (G418 Sulfate, ThermoFisher #10131035) for a week. Fluorescence signals were continuously monitored during the selection process. Upon recovery, G418 at a concentration of 0.5 mg/ml was used for stable cell line maintenance. FACS-sorting for moderate-expression cells was performed to enhance the homogeneity of cells containing labeled mitochondria.

Lentivirus construction of COX8A-mStayGold and Mito-R-GECO1 was constructed with an RSV promoter, EF-1a promoter, and puromycin resistant gene using VectorBuilder custom pipeline. Virus was added at 20 particles per cell (ppc) with 10 µg/ml polybrene (PL200) and incubated overnight. Cells then recovered in normal media for 4-7 days. Selection was performed with 1 µg/ml Puromycin Dihydrochloride (Thermo #A1113803). FACs-sorting was used to filter for high-expression cells (top 10%) to enhance homogeneity and maximize fluorescent intensity signals. Cells infected with both constructs were engineered in sequential transductions.

Dye staining was performed to cells as follows; PKmito Orange was applied at 500 nM for 1 hour at 37°C standard incubation, followed by 2x media wash, two hour incubation, and another 2x media wash. Sybr Gold staining was performed 10,000x dilution for 30 minutes at 37°C standard incubation, followed by a 3x wash. Fluo4 was added 2 µM for 30 minutes at 37°C and then addition of 0.02% Pluronic™ F-127 (Thermo #P3000MP). Calbryte^TM^ 520 AM was added at 1 µM for 1 hour at 37°C followed by a 3x media wash and addition of the anion transporter inhibitor, probenecid (1mM). Fluo4 was added for 30 min at 1 µM followed by a 1x media swap and addition of 0.02% Pluronic™ F-127 (Thermo #P3000MP).

### Live cell imaging

Live cell imaging was performed on several systems including an in-house single-objective light-sheet (SOLS) microscope, a Nikon CSU-W1 spinning disk microscope with laser stimulation, and a Leica Stellaris TauSTED microscope. For all microscopes, cells were plated on fibronectin coated (Sigma #F1141-1mg) 8-well glass-bottom chamber slides (CellVis #C8-1.5H-N) and incubated overnight prior to imaging.

### SNOUTY Light Sheet

The in-house single-objective light-sheet (SOLS) microscope (14) was equipped with a 100x NA 1.35 Silicone primary objective, a fast optical scanner (galvo) for quickly taking 3D data (up to ∼100x100x30µm XYZ field of view), a Gaussian light-sheet (∼12µm Rayleigh range) and a stage-top incubator for maintaining a temperature of 37°C with 5% CO2 throughout. The majority of the imaging was acquired at a frequency of 1 volume/second (1 ms exposures per image) using 5% 488 nm and 12% 561 nm laser powers (20.67 uJ and 70.47 uJ per volume on the sample, respectively) and a quad emission filter (zET405/488/561/635m). Max projected 2D image previews were generated from the 3D data. Deskewing of the 3D light-sheet data was computed using Snouty Viewer (v0.2.5).

### CSU-W1 Spinning Disk and FRAP Laser-stimulation

Laser stimulation was performed on Nikon CSU-W1 spinning-disk confocal system paired with Andor Zyla sCMOS camera (5.5 megapixels), Nikon Perfect Focus system, Oko stage top incubator, and Vortran 405 nm photoactivation and photobleaching laser. Imaging was performed with 488 nm and 561 excitation lasers using a Plan Apo VC 100x/1.4 oil objective and a zET405/488/561/635m quad filter. FRAP laser was calibrated before each experiment using a 0.1mg/ml fluorescein-coated coverslip (Thermo #L13251.36). Laser power was recorded with a Microscope Power Slide Meter Sensor (ThorLabs #S175C). Full 3D z-stacks were acquired with 250 nm slices for a total of 5.25 µm depth across the cell. Imaging 2D time series videos were collected at 0.33 frames sec-1. For autofluorescence detection the camera was swapped to Andor DU-888 EMCCD camera ideal for amplifying low-emission signals.

### Stimulated Emission Depletion Microscopy

For cristae labeling, cells were incubated with 500nM PKmito orange (PKMO, #CY-SC053, Cytoskeleton Inc. (15) for 1 hour at 37°C, followed by 4 successive washes over a two hour period. Additionally, mtDNA nucleoid staining was performed with 1:10,000x SybrGold (#S11494, ThermoFisher) for 30 min incubation at 37°C followed by a 3x wash. Images were acquired using an 86x water objective, taken at 7x zoom, 16x line averaging, and 1024x524 resolution with PKMO acquired with a 5% 591 nm white light laser excitation line, a 70% 775 nm STED laser depletion line, and SybrGold at 4% 488 nm excitation. Tau-Sted lifetime enhancement was applied at 100%. Pearling induced with 405 nm stimulation laser fired at 20 mW for 200 milliseconds.

### Focused Ion Beam Scanning Electron Microscopy (FIB-SEM)

Whole-cell FIB-SEM data was obtained with author’s permission from the Janelia OpenOrganelle Platform using samples ‘jrc_ctl-id8-2’, ‘jrc_ctl-id8-3’, and ‘jrc_ctl-id8-4’, previously published with Ritter et al.(16) Briefly, OT-1 mouse cytotoxic T-cells were isolated and activated before combining with adherent ID8 cancer cells (ATCC) on a sapphire coverslip (corning). ID8 and T-cells cells were labeled with mEmerald-Chmp4b and TOMM20-SNAP, and the media was loaded with propidium iodide to detect membrane puncture of the target cell. Samples were high-pressure frozen and cryogenic fluorescent imaging was used to select cells. Cells were selected for presence of a propidium iodide gradient signature at the synapse between spatially adjacent T-cells and ID8 cells. After resin processing, a customized FIB-SEM with a Zeiss Capella FIB column mounted at 90° onto a Zeiss Merlin SEM imaged with an 8-nm x,y pixel size followed by repetitive 4nm gallium beam ablations. Image registration and alignment was performed with a Scale Invariant FeatureTransform (SIFT) algorithm based Fiji plug-in(23). Every two consecutive images were binned and averaged down to one to obtain 8x8x8 nm^3^ isotropic voxels. Mitochondrial segmentation was performed with the 3dEMtrace platform of ariadne.ai (https://ariadne.ai/) and custom thresholding to isolate membrane layers. Cristae 3D reconstructions used an Skimage erosion function to remove the inner membrane surface layer, followed by rendering in the Napari GUI. For complete methods see Ritter et al.(16)

### Micro-needle manipulation

Microneedle manipulation was adapted for interphase U2OS cells based on previous techniques detailed in Suresh et al. Microneedles were made from glass capillaries with an inner and outer diameter of 0.58 mm and 1 mm respectively (1B100-4 or 1B100F-4, World Precision Instruments). Glass capillaries were pulled via micropipette puller (P-87, Sutter Instruments, Novato, CA), bent and polished using a microforge (Narishige International, Amityville, NY) according to the same specifications, parameters, and geometries described in detail in Suresh et al. These parameters allowed for the needle to approach cells orthogonal to the imaging plane and conduct manipulations without rupturing the cell (Suresh et al.) Prior to imaging microneedles were coated with BSA Alexa Fluor 647 (A34785, Invitrogen) by soaking them in solution for 60s. The solution was created via dissolving BSA-Alexa dye and Sodium Azide (Nacalai Tesque, Kyoto, Japan) in 0.1 M phosphate-buffered saline (PBS) at a final concentration of 0.02% and 3 mM, respectively (Sasaki et al., 2012). The solution allows needles to be visualized via fluorescence imaging, aiding in positioning of the needle along a single mitochondrion. It should be noted, that due to technical constraints involved with manipulation experiments, the imaging chamber lid had to be removed. As such, the sample dish was only able to achieve a stable temperature of 30 C.

Cells for microneedle manipulation were chosen based on the following criteria: chosen cell was in interphase and cell possessed one or more mitochondria that are well separated from the surrounding mitochondria network. These criteria were important for perturbing single mitochondrion without affecting the entire network.

The micromanipulator was mounted to the scope body and positioned above samples as described in Suresh et al. Manipulations were performed in 3D using an x-y-z stepper-motor micromanipulator (MP-225, Sutter Instruments, Novato, CA). A 3-axis-knob (ROE-200) connected to the manipulator via a controller box (MPC-200, Sutter Instruments). Prior to manipulation, the needle was positioned via phase imaging at the approximate x-y position of the intended manipulation site ∼ 10 μm above the cell in z. While imaging every 2 seconds at 3 z-planes (0.0 +/- 0.5 μm) the needle was manually lowered into place near a single mitochondrion. If necessary, the needle’s position in x-y was changed by first raising the needle above the cell and repositioning in x-y before approaching the intended mitochondrion. Once properly positioned, the needle was manually moved to tap the mitochondrion to apply a brief mechanical load. After, the needle was slowly removed from the cell.

To be analyzed, the manipulated cells had to demonstrate all listed attributes: cell health was not significantly impacted via manipulation (ie cell rupture), mechanically challenged mitochondrion was clearly visible throughout the entirety of manipulation, the mitochondrion demonstrated slight deformation due to the applied mechanical load, a single mitochondrion was contacted by the needle.

### Image Analysis

Image analysis was performed in fiji software as well as an updated version of MitoMeter (17) called Nellie (18) and available at https://github.com/aelefebv/nellie. This pipeline enables automated segmentation, tracking, and feature extraction of individual organelles over time. Metrics derived from Nellie include mitochondrial counts, tubule length, tubule width, aspect ratio, linear velocity, linear acceleration, angular velocity, angular acceleration. A separate pearl detection script was added on the end of Nellie specifically to identify pearls and measure pearl shape. First 3D volumes were max projected into 2D, and a polygon mask was drawn over the pearled region. Pearls were then detected by applying Frangi filtering to enhance tubular structure within each image. The filtered images were thresholded to binary masks, followed by KDtree-based distance transformation and maximum filtering to identify local maxima for potential pearls. Candidate peaks were labeled and refined using a proximity-based region growing algorithm, followed by removal of small objects, and those touching image edges to eliminate false positives.

### Frequency of pearling events

The frequency of pearling events was calculated with two approaches. The first method counted the number of pearling events in each video and normalized by the duration of imaging. The total number of mitochondria was measured using the Nellie software object detection, yielding the number of events per mitochondria per second. The event probability over 10 minutes was then calculated. The second method measured the total length of pearled mitochondria across a minute of time, and divided by the sum of lengths across all mitochondria in the cell outputted by Nellie software. To distinguish pearling from other dynamics such as fission events, inclusion criteria for pearling included, i) the mitochondria underwent a full transition from cylinder to pearls to cylinder. ii) the pearl formation contained at least three pearls, and iii) the pearls remained connected by visible fluorescent signal between one another during the transition.

### Intensity-based curvature

Curvature metrics were calculated on tracked skeleton line profiles of individual mitochondrial tubules over time outputted by Nellie software. Curvature was calculated using a 2D parametric curve formula to a spline fitted line across the skeleton. Smoothing factors were optimized for each time series to maximize the difference in curvature between a pearled morphology and the frame before the onset of pearling. Curvature was weighed by the length of the skeleton and normalized to the first 3 timepoints before the onset of pearling. Other extracted metrics include peak-to-peak distance, volume, length, width, height, duration of event, pre/post tortuosity, the duration of the full transition, time to peak of event, time of recovery, number of pearls, length of pearled segment, pearls per micron, width/length of tubule before and after pearling, number of pearling events per tubule.

## Supplemental Tables

**Table S1.**
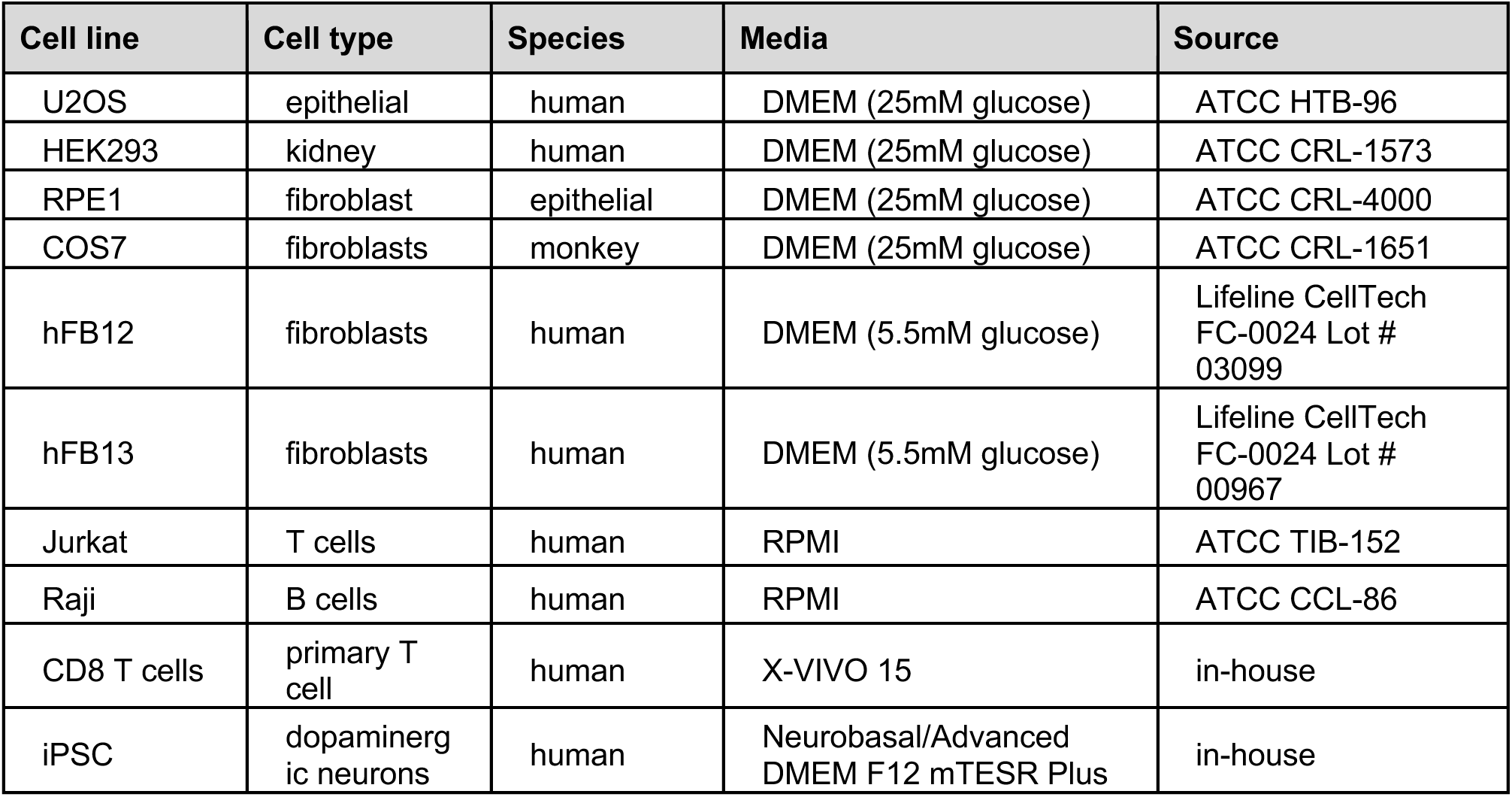
List of mammalian cell types images for mitochondrial pearling dynamics.

| Cell line | Cell type | Species | Media | Source |
| --- | --- | --- | --- | --- |
| U2OS | epithelial | human | DMEM (25mM glucose) | ATCC HTB-96 |
| HEK293 | kidney | human | DMEM (25mM glucose) | ATCC CRL-1573 |
| RPE1 | fibroblast | epithelial | DMEM (25mM glucose) | ATCC CRL-4000 |
| COS7 | fibroblasts | monkey | DMEM (25mM glucose) | ATCC CRL-1651 |
| hFB12 | fibroblasts | human | DMEM (5.5mM glucose) | Lifeline CellTech<br>FC-0024 Lot #<br>03099 |
| hFB13 | fibroblasts | human | DMEM (5.5mM glucose) | Lifeline CellTech<br>FC-0024 Lot #<br>00967 |
| Jurkat | T cells | human | RPMI | ATCC TIB-152 |
| Raji | B cells | human | RPMI | ATCC CCL-86 |
| CD8 T cells | primary T cell | human | X-VIVO 15 | in-house |
| iPSC | dopaminergic neurons | human | Neurobasal/Advanced<br>DMEM F12 mTESR Plus | in-house |

**Table S2.** Yeast knockout strains screened for pearling.

| Knockout | Yeast strain | Target | Label |
| --- | --- | --- | --- |
| HO<br>(Control) | CGY14.31xYKO<br><ho> | HO | preCox4 mNeon-green |
| CRD1 | CGY14.31xYKO<br><crd1> | cardiolipin synthesis | preCox4 mNeon-green |
| DNM1 | CGY14.31xYKO<br><dnm1> | fission machinery | preCox4 mNeon-green |
| FIS1 | CGY14.31xYKO<br><fis1> | fission synthesis | preCox4 mNeon-green |
| TAZ1 | CGY14.31xYKO<br><taz1> | cardiolipin synthesis | preCox4 mNeon-green |
| CRD1 | CGY14.31xYKO<br><crd1> | cardiolipin synthesis | preCox4 mNeon-green |
| GEM1 | CGY14.31xYKO<br><gem1> | ERMES complex | preCox4 mNeon-green |
| Control | CGY49.77 | wildtype | preCox4 mNeon-green |
| MIC10 | CGY49.77xYKO<br><mic10> | MICOS complex subunit | preCox4 mNeon-green |
| MIC12 | CGY49.77xYKO<br><mic11> | MICOS complex subunit | preCox4 mNeon-green |
| MIC19 | CGY49.77xYKO<br><mic19> | MICOS complex subunit | preCox4 mNeon-green |
| MIC60 | CGY49.77xYKO<br><mic60> | MICOS complex subunit 60 | preCox4 mNeon-green |

**Table S3:** IMM = inner mitochondrial membrane.

| Label | Category | Target | Properties | Source |
| --- | --- | --- | --- | --- |
| MitoTracker Green FM | dye | mitochondrial matrix | membrane potential dependent | Thermo #M7514 |
| PKmito Orange | dye | IMM | membrane potential dependent | Cytoskeleton Inc #CY-SC053 |
| Sybr Gold | dye | mtDNA intercalation | membrane potential dependent (partial) | Thermo #S11494 |
| TMRE | dye | mitochondrial inner matrix | Mitochondrial membrane potential dependent | Thermo # T669 |
| CellLight Mito-GFP | transient plasmid | PDHA1 IMM and matrix | Marker of mitochondrial structure | Thermo #C10600 |
| CellLight Mito-RFP | transient plasmid | PDHA1 IMM and matrix | Marker of mitochondrial structure | Thermo #C10601 |
| CellLight ER-RFP | transient plasmid transfection | ER | Marker of ER structure | Thermo #C10591 |
| COX8A-mEmerald | genetically encoded sequence | COX8A protein on the IMM and mitochondrial matrix | geneticin selection, FACS-sorted | Addgene #54160 |
| COX8A-mStayGold | lenti-transduction | COX8A protein on the IMM and mitochondrial matrix | lentivirus package, puromycin selection, FACS-sorted | https://doi.org/10.1038/s41592-023-02085-6 |
| Mito-R-GECO1 | lenti-transduction | mitochondrial calcium | ratiometric intensity mitochondrial calcium flux | Addgene #46021 |
| MitoSOX red | dye | mitochondrial ROS | mitochondrial superoxide indicator | Thermo #M36008 |
| MitoSOX green | dye | mitochondrial ROS | Superoxide indicator | Thermo #M36005 |
| Hyper7-IMS | dye | mitochondrial H<sub>2</sub>O<sub>2</sub> ROS | mitochondrial intermembrane space hydrogen peroxide indicator | Addgene #136469 |
| Calbryte™ 520 AM | dye | intracellular calcium | cytoplasmic calcium flux | AAT Bioquest #20651 |
| Mito Flipper-TR | dye | mitochondrial membrane tension | mitochondrial membrane tension | Cytoskeleton #CY-SC023 |

**Table S4.** Chemical inhibitors of mitochondria.

| Inhibitor | [ ]<br>(μM) | Duration<br>(min) | Known target | Pearling<br>effect | Source |
| --- | --- | --- | --- | --- | --- |
| FCCP | 10 | 10-60 | mitochondrial membrane potential | full inhibition | Sigma #C2920 |
| Oligomycin | 1 | 10-60 | ATP synthase inhibitor | inhibits recovery | Sigma #75351 |
| Rotenone | 4 | 10-60 | complex II inhibitor | partial inhibition | Sigma #R8875 |
| Antimycin A | 2 | 10-60 | complex II inhibitor | partial inhibition | Sigma #A8674 |
| Latrunculin B | 10 | 30-60 | actin polymerization | inhibits recovery | Sigma #L5288 |
| Nocodazole | 10 | 30-60 | microtubule polymerization | partial inhibition | Sigma #M1404-2MG |
| Taxol | 10 | 30-60 | microtubule stabilization | inhibits recovery | Sigma #PHL89806 |
| Cyclosporin A | 10 | 10-60 | mPTP inhibitor | increased | Sigma #30024 |
| Atorvastatin | 50 | 240 | cholesterol depletion | full inhibition | Selleck Chemical # S207750MG |
| NAC | 5 | 24 hour | cytoplasmic antioxidant | n.s. | Thermo #A15409.36 |
| MitoTEMPO | 0.02 | 24 hour | mitochondrial-specific antioxidant | n.s. | Sigma #SML0737 |

## Supplemental Figures

**Figure S1.**
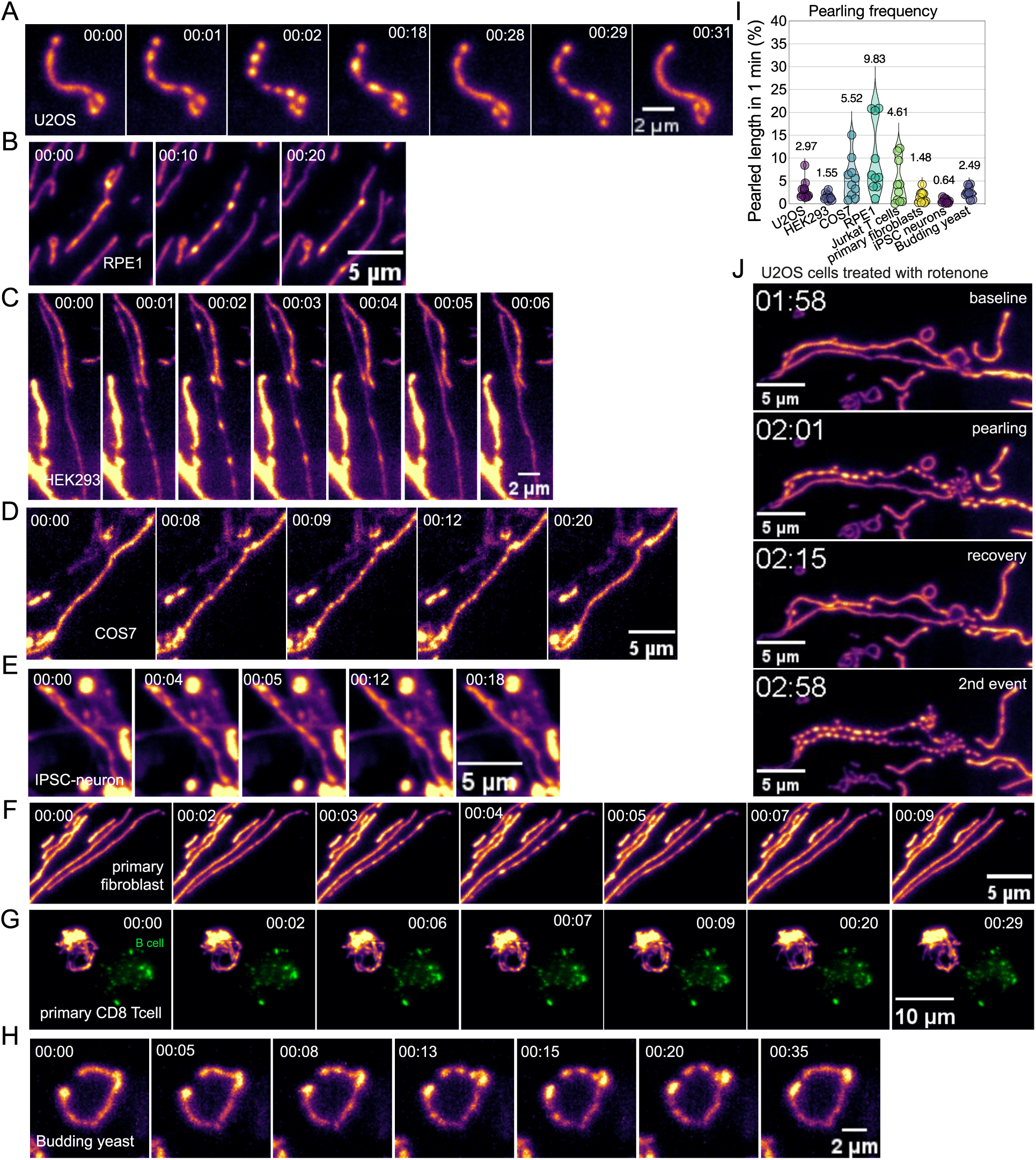
Panel of pearling by cell type. (**A-H**) Transient mitochondrial pearling occurring spontaneously under light sheet microscopy (SNOUTY), imaging at 1 3D volume per second across various cell types. HEK293, RPE1, COS7, primary fibroblast, iPSC neuron, and primary CD8 T cell labeled with PKmito Orange (IMM), U2OS labeled with COX8a-mStayGold (IMM). The green cell in the bottom left corner in the primary T cell video is a Raji cell labeled with SybrGold and pre-treated with the superantigen SEE to induce immune synapse formation. (**I**) Rate of pearling per µm of length measured as the percent of mitochondrial length that has pearl within one minute period (n=10-11 videos per group). (**J**) Spontaneous pearling of mitochondria tubule of a U2OS cell. U2OS cells treated with rotenone (5 µM for 30min) undergo a pearling transition twice. Mitochondrial inner membrane (inferno LUT) labeled with genetically-encoded COX8A-mEmerald.

**Figure S2.**
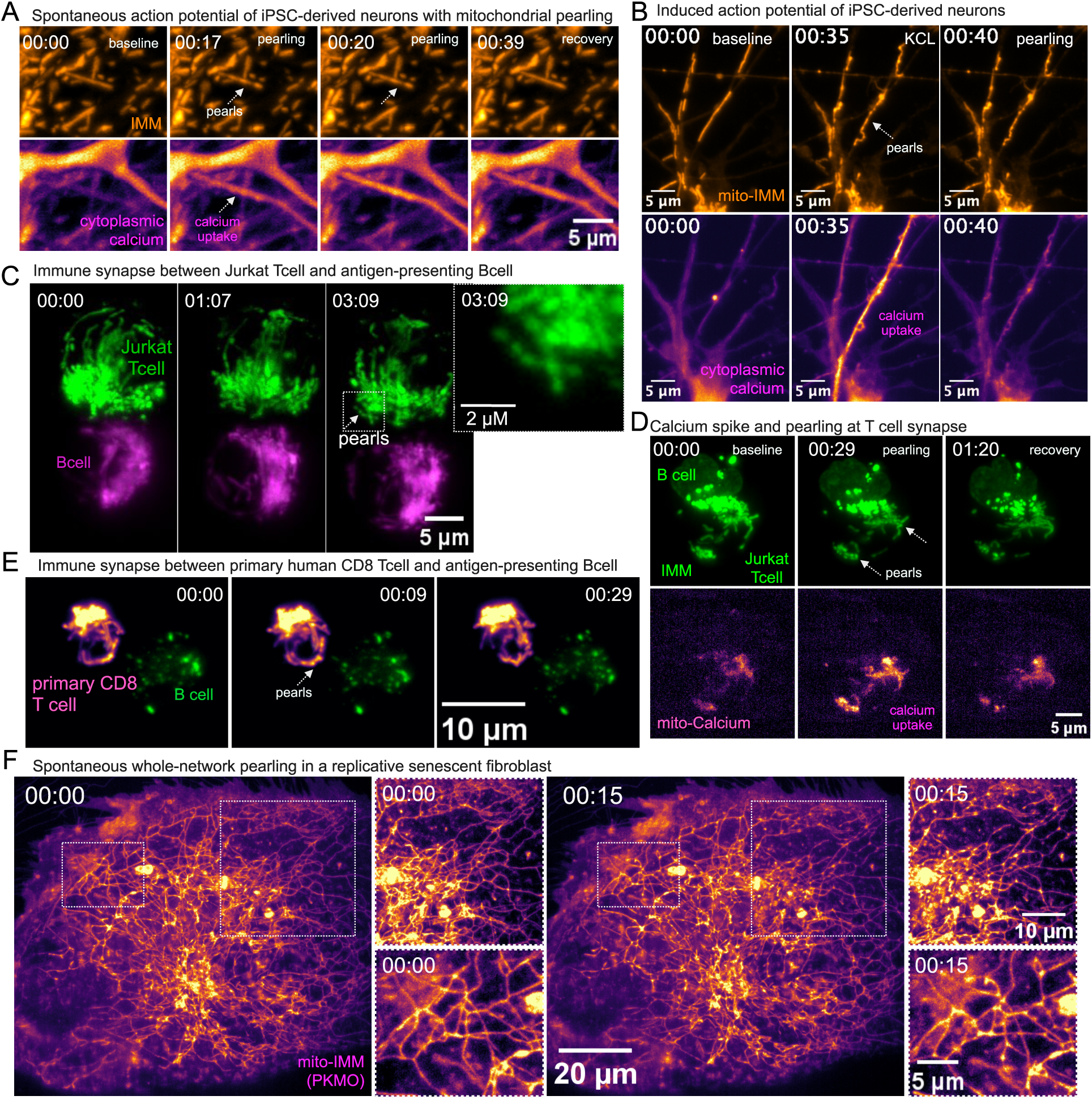
Mitochondrial pearling observed across physiological events marked by ionic flux. (**A**) Spontaneous mitochondrial pearling coinciding with a spontaneous action potential in an iPSC-derived neuron. Mitochondrial inner membrane labeled with PKmito Orange. Cytoplasmic calcium labeled with CalBryte^TM^ 520am. (**B**) Pearling observed during a KCL (50 mM) triggered action potential of iPSC-derived dopaminergic neurons. Top panel tracks the mitochondrial inner membrane using the dye PKmito Orange. Bottom-panel tracks global calcium levels using Calbryte^TM^ 520 AM dye. (**C**) Curvature per µm over time normalized to the first three time points. Dotted line indicates the point of 50mM KCl injection (n=5 cells). (**D**) Mitochondrial pearling observed at the synapse between a Jurkat cell labeled with Mito-mStayGold targeting an SEE-pulsed Raji cell labeled with PKmito Orange. Both labels mark the mitochondrial inner membrane. (**E**) Mitochondrial pearling observed at the synapse between a primary human CD8+ T cell labeled with PKmito orange (IMM) targeting an SEE-pulsed Raji cell labeled with Sybr Gold (mtDNA). (**F**) Spontaneous pearling observed across an entire hyper-fused mitochondria network of a replicative senescent primary human fibroblast. Rapid retraction events are visible.

**Figure S3.**
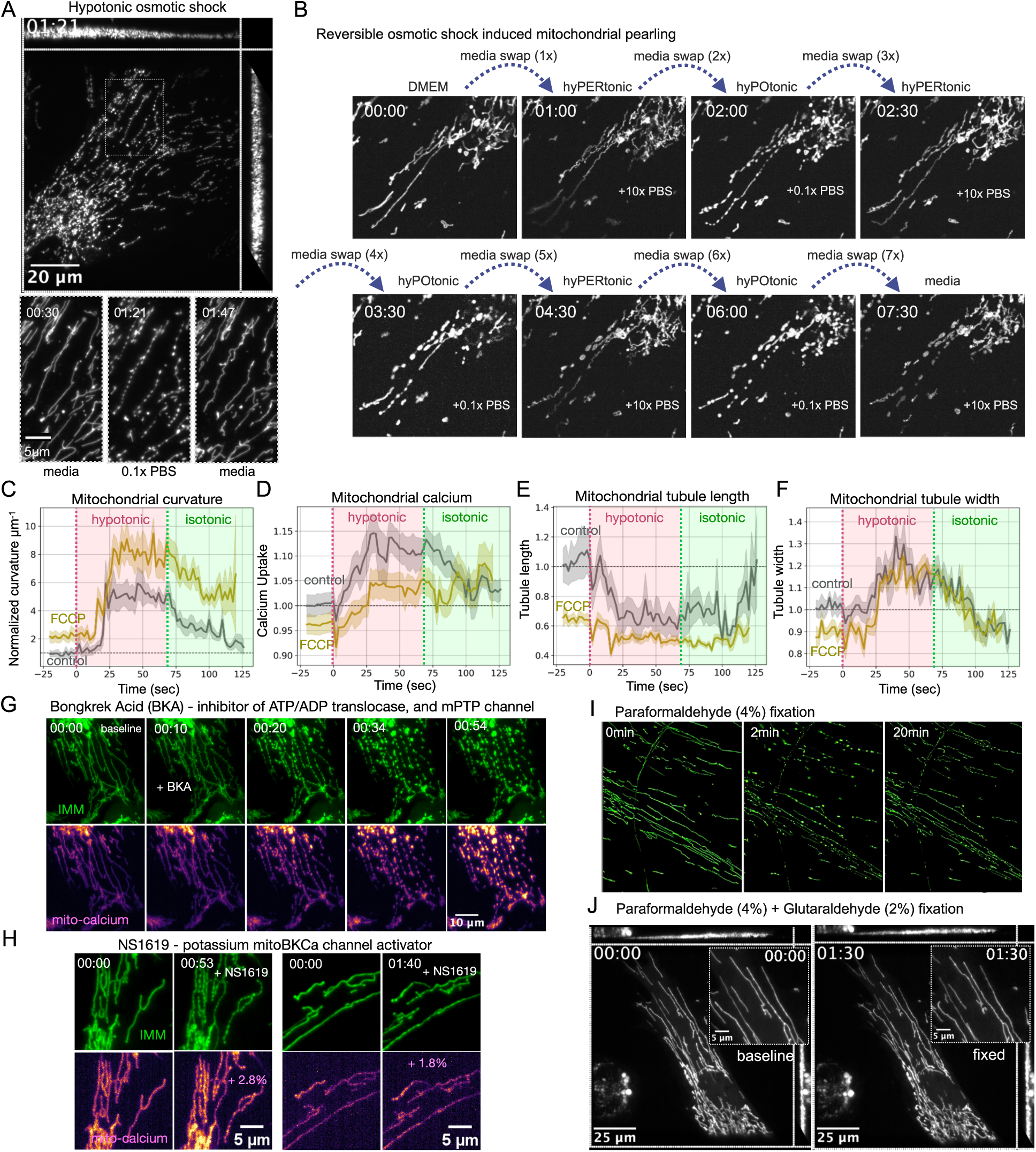
Osmotic shock and chemical induction of mitochondrial pearling. (**A**) Mitochondrial pearling induced by replacing DMEM with hypotonic medium, 0.1x PBS and then recovered by replacing medium back to DMEM. (**B**) Reproduction of the 1915 experiment (Lewis & Lewis, AJA, 1915) showing reversible pearling morphology via altering from hypo-osmotic (0.1x PBS) to hyper-osmotic (10x PBS) media conditions. The mitochondria of human epithelial U2OS cells are marked with genetically encoded COX8A-mEmerald. (**C-F**) Time series of untreated (gray) and cells pre-treated with 10 µM FCCP for 15min (yellow) with (F) Normalized curvature per µm (COX8a-mStayGold), (G) mitochondrial calcium uptake (Mito-R-GECO1) and (H) mitochondrial tubule length (COX8a-mStayGold). n = 12 cells per group. Data normalized the mean of the first 5 control timepoints. Red vertical dotted line indicates media swap to hypotonic shock (0.1x PBS). Green line indicates media swap to isotonic DMEM. (**G**) Time series of 10 µM BKA induced pearling with a transient ‘hypertonic’ signature visible at 10 seconds, which precedes the rise in calcium. Top panel is labeled with IMM (COX8A-mStayGold), bottom-panel is labeled with mitochondrial calcium (Mito-R-GECO1). (**H**) Time series of the potassium BKCa channel activator NS1619 (10 µM) induced pearling with or without calcium uptake. Top panel is labeled with IMM (COX8A-mStayGold), bottom-panel is labeled with mitochondrial calcium (Mito-R-GECO1). (**I**) Pearling induced by fixation with 4% paraformaldehyde (PFA) and remains pearled indefinitely. (**J**) Stable mitochondrial network in primary human fibroblast fixed by combining 4% PFA with 2% glutaraldehyde (GA).

**Figure S4.**
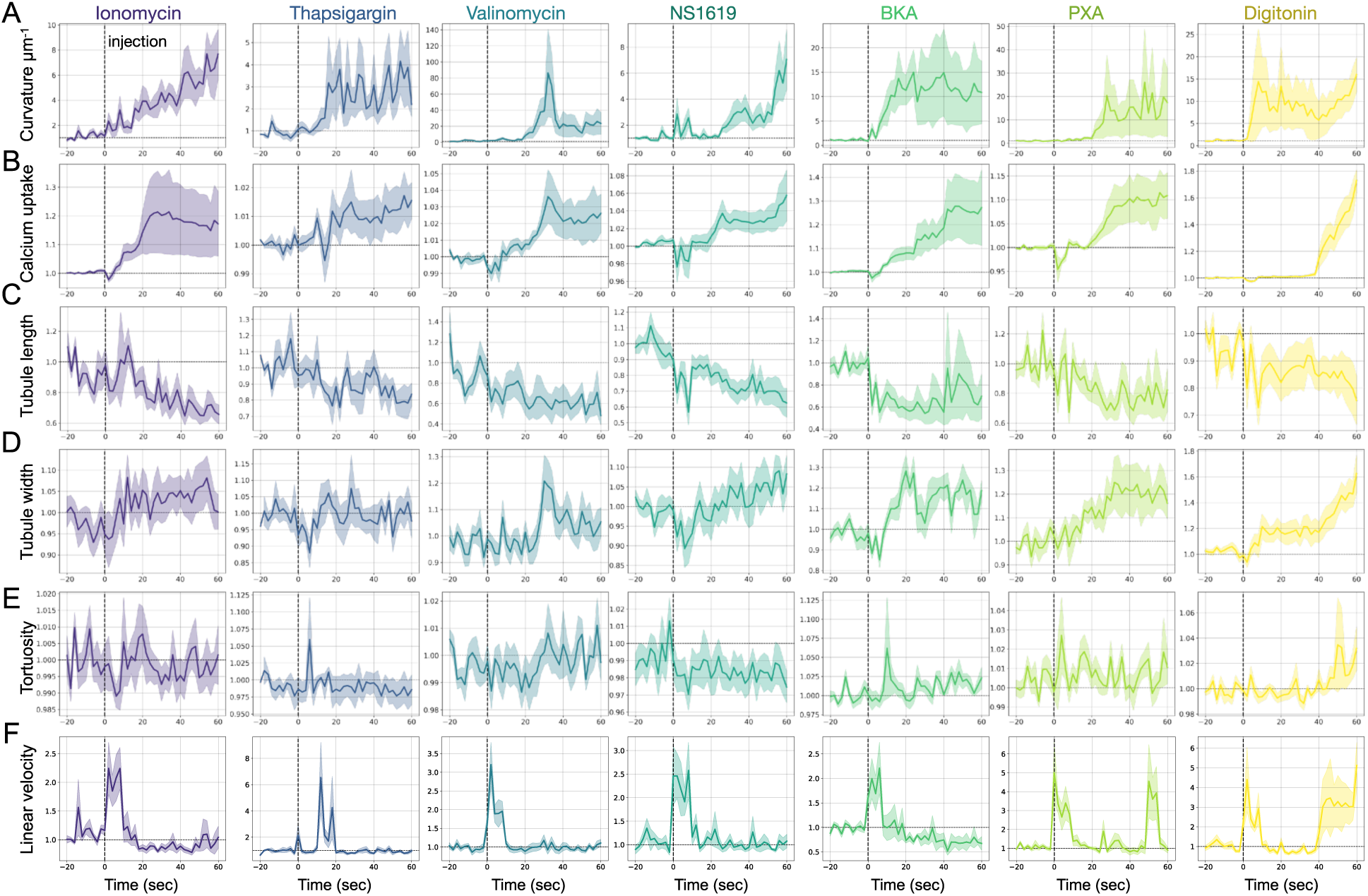
Dynamical metrics tracked after chemically induced pearling. (**A-F**) Tracked feature of mitochondrial pearling induced, from left to right, by ionomycin (4 µM), thapsigargin (10 µM), valinomycin (10 µM), NS1619 (10 µM), BKA (10 µM), PXA (10 µM), and digitonin (10 µM). All data is normalized to the average of the first 5 timepoints of that treatment. Tracked metrics include; (**A**) normalized curvature per µm measured on IMM marker COX8A-MitoStayGold, (**B**) calcium uptake as measured by the change in mean fluorescence intensity of Mito-R-GECO1, (**C**) mean tubule length, (**D**) mean tubule width, (**E**) tortuosity (**F**) linear velocity (µm^2^). Black dotted line indicates the time at which the chemical was added.

**Figure S5.**
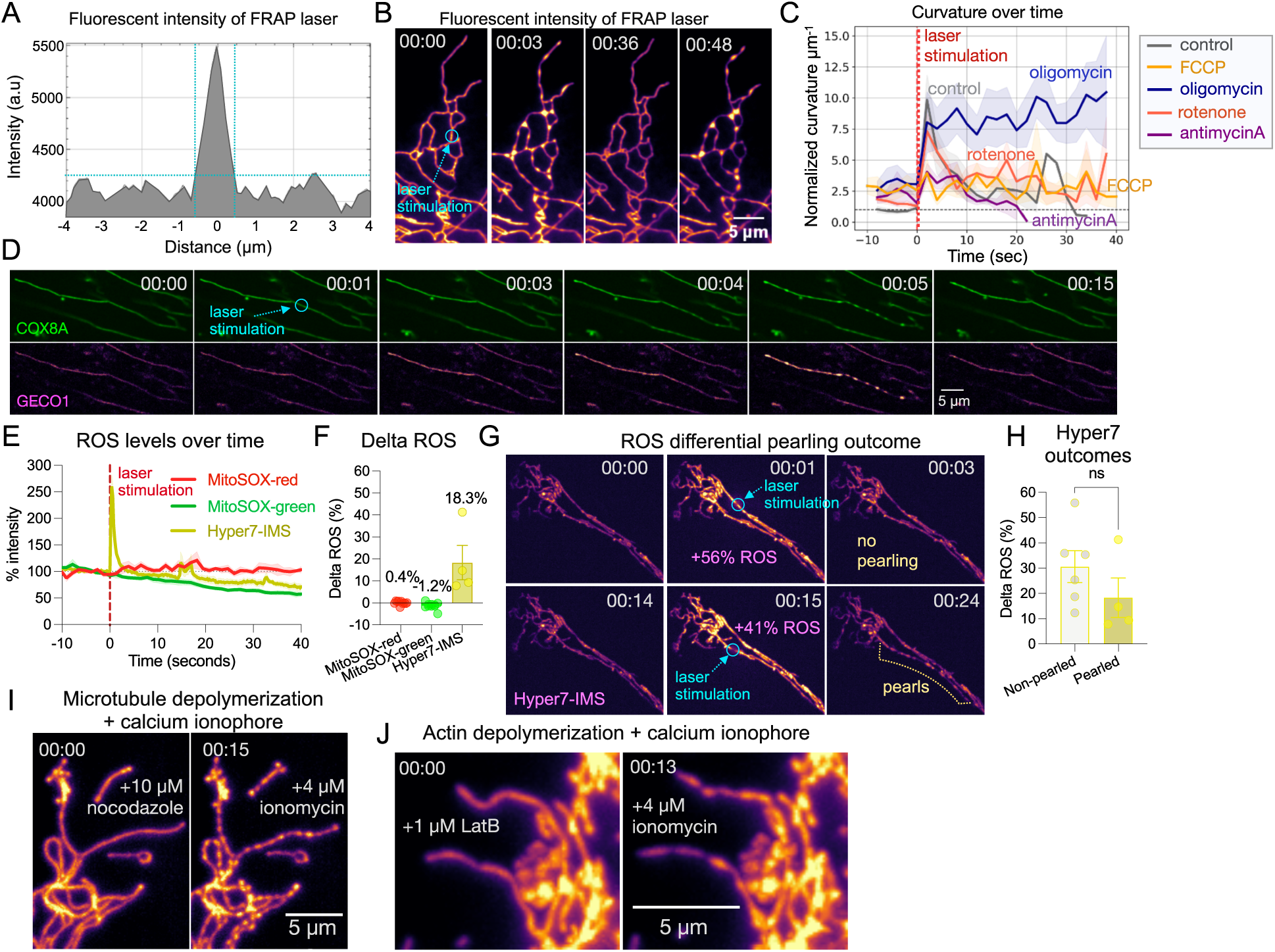
Characterizing laser-induced pearling events. **(A**) Precision of 405nm stimulation laser, Crest-C2, with 100x oil immersion objective, 20mW (0.3mW reaching sample) for 200 milliseconds. (**B**) Laser-induced pearling recovers back to cylindrical shape but then spontaneously returns back to the pearled shaped. (**C**) Curvature per unit µm comparing electron transport chain inhibitors, n=21 oligomycin (1 uM) events, n=18 FCCP (4 µM) events, n=18 rotenone (5 uM) events, n=18 antimycin A (5 uM) events. Red dotted line indicates point of laser stimulation. All data normalized to the average of the first five timepoints of the control group. (**D**) High-temporal tracking of calcium rise that precedes the onset of laser-induced pearling. COX8A = IMM and GECO1 = mitochondrial calcium. (**E**) ROS fluorescent sensors tracked over time after laser-induced pearling event (n=10 events per sensor). Data normalized to the average of the 5 timepoints before stimulation. (**F**) ROS sensors percent change in fluorescent intensity after laser-stimulation that induced pearling events. (**G**) Representative example of Hyper7-IMS ROS sensor increasing across mitochondrial network after laser stimulation, regardless of pearling event occurring. (**H**) Hyper7-IMS percent change in fluorescent intensity after laser-stimulation for tubules that remained uniform (left bar) and those that underwent a pearling event (right bar). Mann-Whitney non-parametric t-test performed. (**I-J**) U2OS cell treated with 10 µM of nocodazole (I) or 1 µM latrunculin B (J) for 1 hour and mitochondria labeled with COX8A-mStayGold (IMM). Pearling is then induced by injection of 4 µM ionomycin.

**Figure S6.**
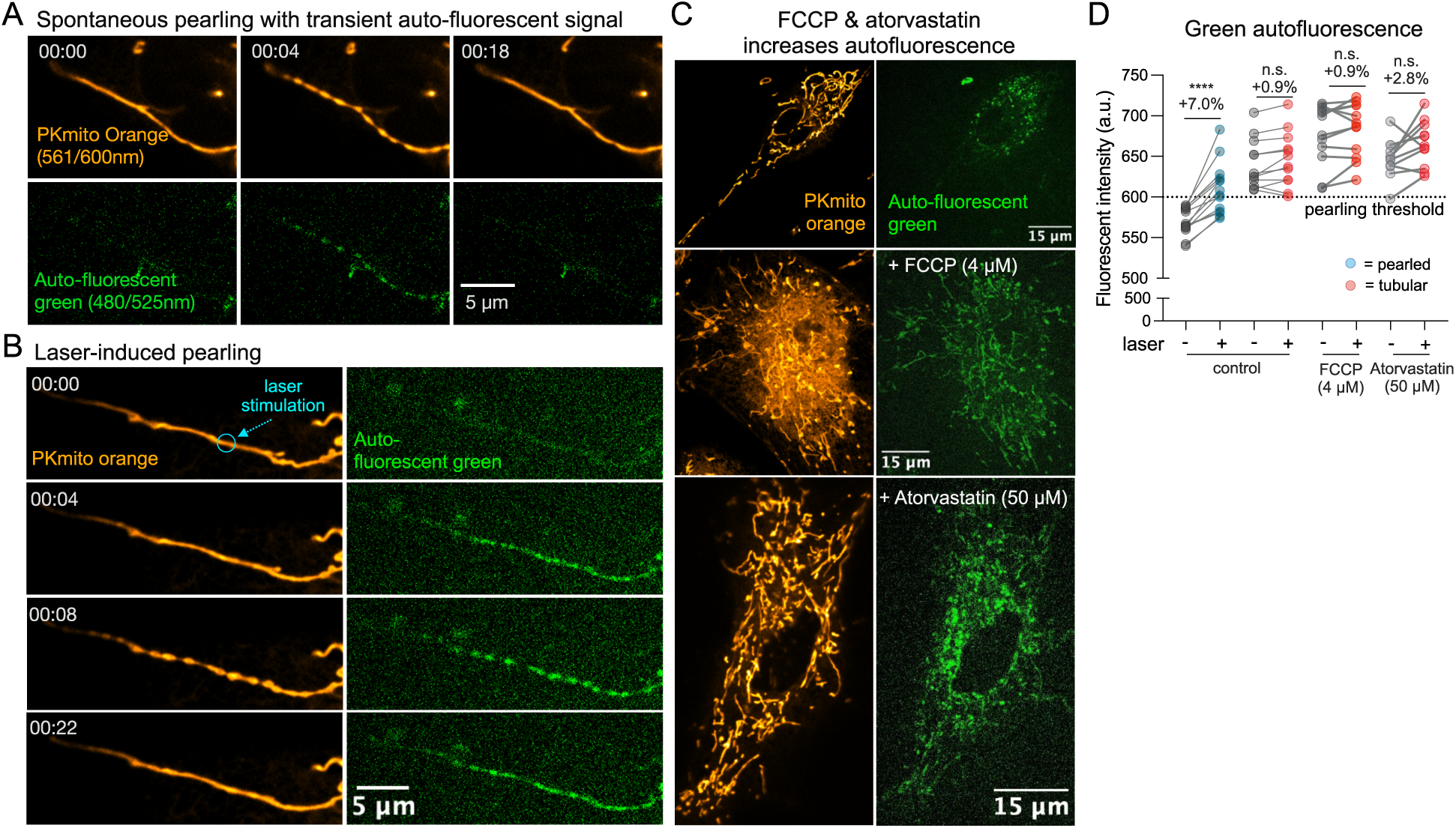
Green autofluorescence predicts mitochondrial pearling transition. (**A**) Spontaneous pearling event is followed by a rise in green-shifted autofluorescence that recovers back to background as the tubule reforms. (**B**) Laser-induced pearling similarly results in a rise in green-shifted autofluorescence within the mitochondrial tubule. (**C**) Cells treated with FCCP (4 µM) and atorvastatin (50 µM) increase green-shifted autofluorescence and do not respond to the laser but rather remain in the tubular state. (**D**) Level of green-shifted autofluorescence within the mitochondrial tubule before and after firing the laser. Mitochondria are split left to right by those that pearl (blue-dots) and those that remain in the tubular state (red-dots). Dotted-line indicates fluorescent threshold at which mitochondria will pearl after firing a 405nm stimulation laser. Mixed-effects anova performed with multiple comparison. **** = 0.00001, n.s. = 0.10 p value.

**Figure S7.**
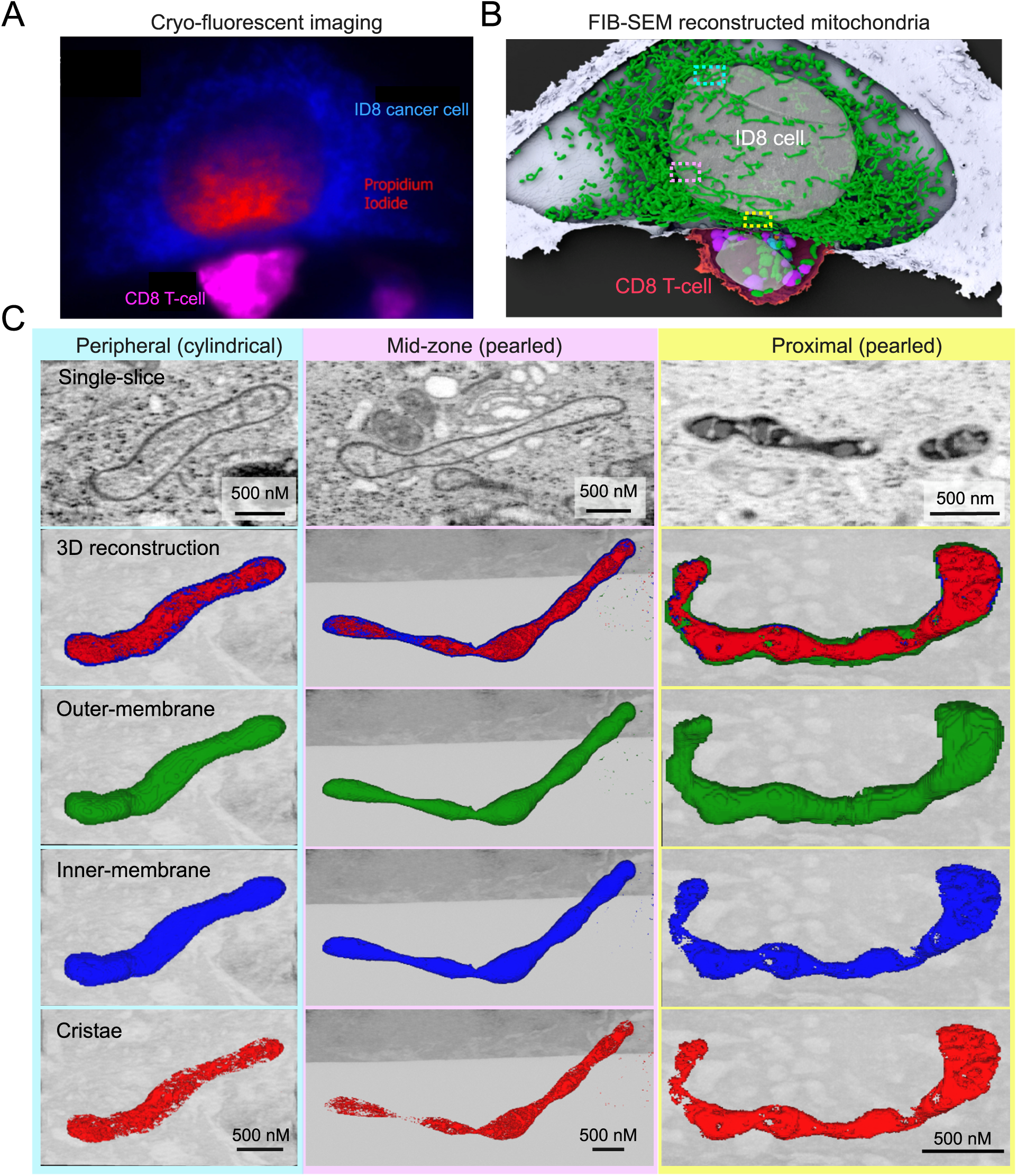
**Pearled mitochondria ultrastructure following cytotoxic T-cell synapse formation** (**A**) Cryo-fluorescent imaging of high-pressure frozen coverslip with selected synapse formation between OT-I mouse cytotoxic T-cell (pink) and ID8 ovarian cancer blue. Cell was selected for scanning electron microscopy with presence of extracellular propidium iodide around the synapse indicating onset of cytotoxic T cell induced cell death. (**B**) FIB-SEM reconstructed mitochondria (green) after the release of perforin and granzyme from lytic granules at the synapse of ID8 cancer cell targeted by a cytotoxic T-cell. Dotted boxes mark selected representative mitochondria in the target cell organized by proximity to the synapse of the CD8 T-cell. (**C**) Representative reconstruction of mitochondrial ultrastructure at the periphery (left-panel), mid-zone (central panel), and proximal (right-panel) to the T-cell synapse. Data and images adapted from Ritter et al. *Science*, 2022 with author permission.

**Figure S8.**
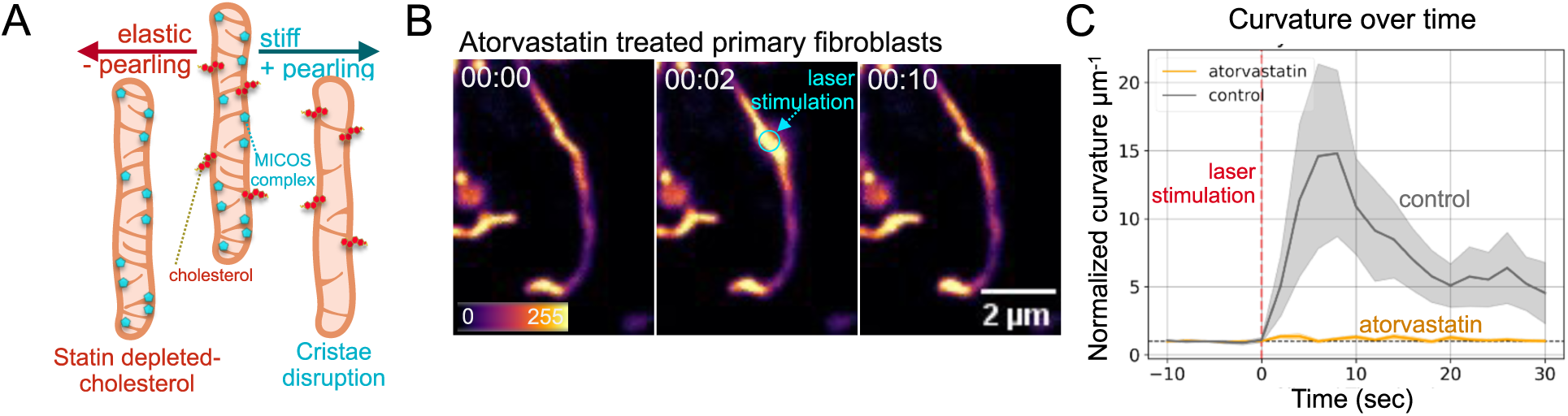
Cholesterol depletion with statins inhibits laser induced pearling. (**A**) Schematic of how membrane content affects the biophysical properties of the mitochondria and influence its likelihood to pearl. (**B**) Mitochondria of a primary fibroblasts treated overnight with 50 µM atorvastatin before and after FRAP laser stimulation (blue circle). Cells are labeled with PKmito Orange. (**C**) Timecourse of the membrane curvature along the mitochondrial tubule of control (n=11) and overnight atorvastatin-treated (24hr of 50µM, n=13) primary human fibroblasts.

**Figure S9.**
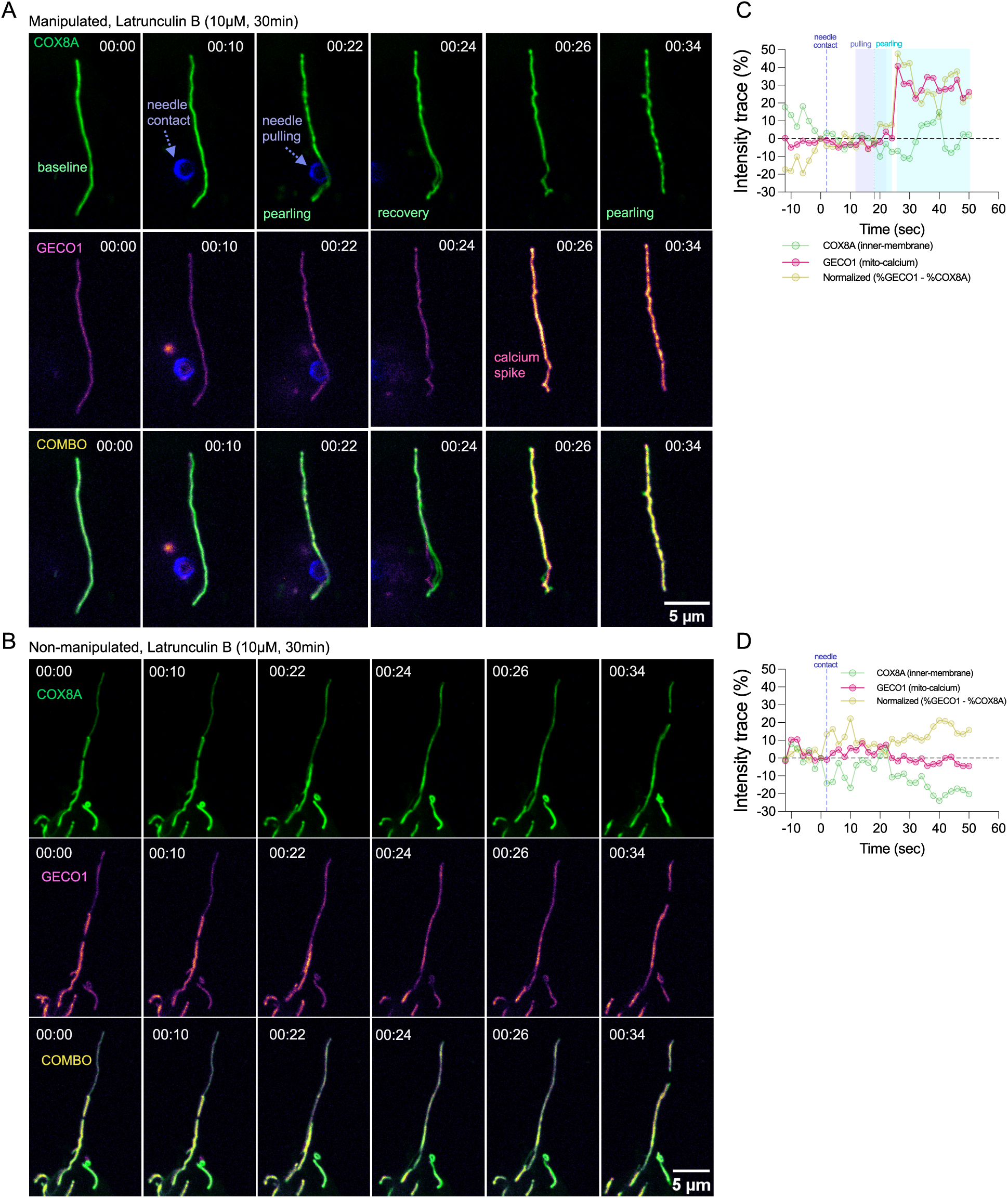
Micro-needle induced pearling precedes calcium uptake. (**A**) Glass micro-needle manipulation of mitochondria in a U2OS cell treated with 1 µm latrunculin B for 30 minutes. (**B**) A tubule from the same cell that was not manipulated by the needle. Cells are labeled with COX8A-mStayGold (inner-membrane) and Mito-R-GECO1 (**C-D**) Intensity trace (R1/R0) over time normalized to the point prior to the presence of the needle in the cell for the (C) manipulated tubule of A and (D) a non-manipulated tubule of B.

**Figure S10.**
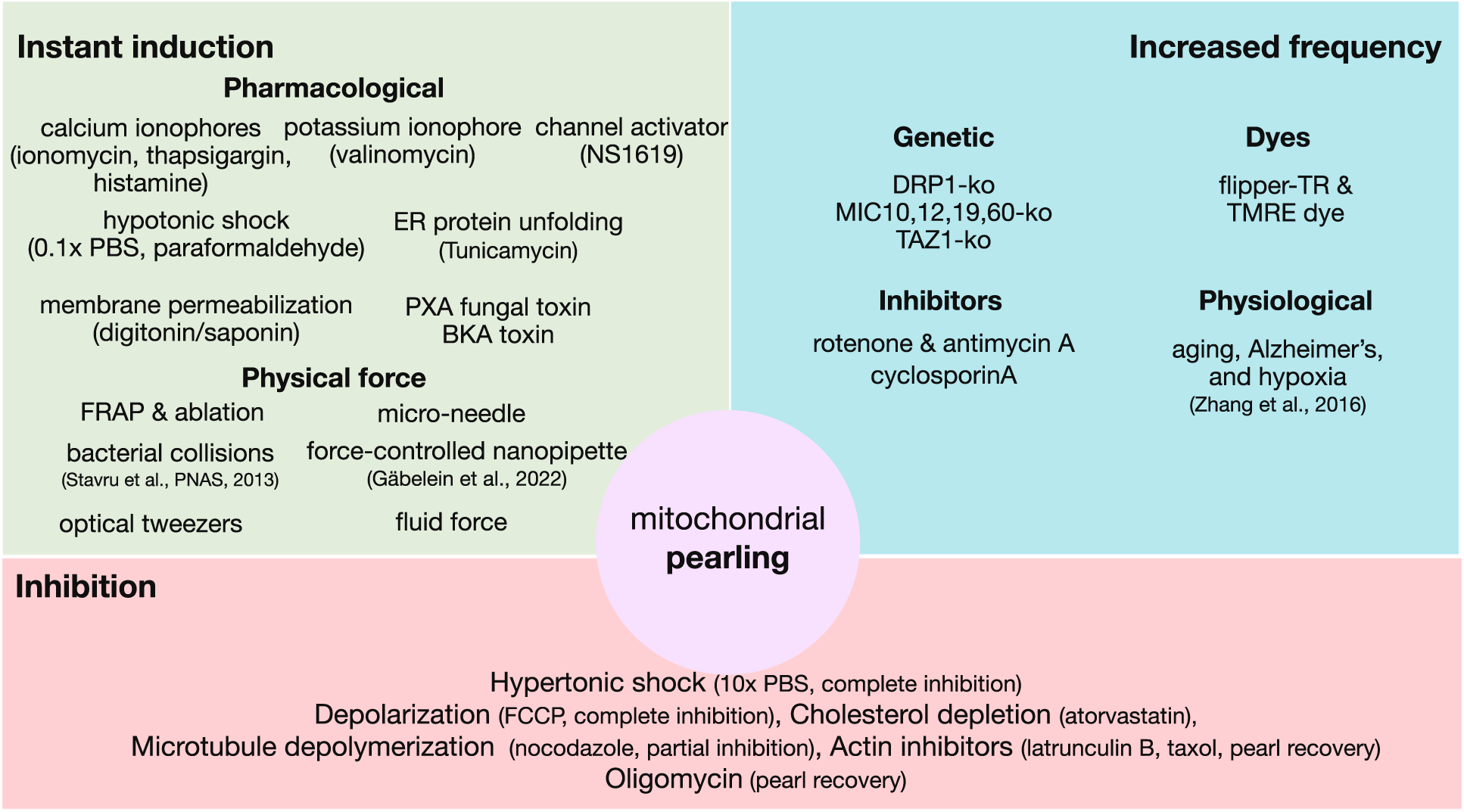
Experimental perturbations that influence mitochondrial pearling. Perturbations are characterized by those that induce pearling across the entire mitochondrial network in the cell (light-green box), those that increase the frequency of spontaneous pearling events (light-blue box), and those that inhibit either the induction or recovery of pearling events (light-red box).

## Supplemental Movie Legends

Movies S1-S38 are available on our public repository at https://github.com/gav-sturm/Mitochondrial_Pearling/tree/main/Supplementary_Movies.

**Movie S1 (Movie01_spontaneous_U2OS_pearling.gif)**

Time-lapse of spontaneous mitochondrial pearling in a U2OS cell labeled with COX8a-mEmerald (IMM). Images taken on SNOUTY at 1 volume/second.

**Movie S2 (Movie02_all_cell_types_spon_pearling.gif)**

Comparative view of spontaneous pearling events across multiple cell types. HEK293, RPE1, COS7, primary fibroblast, iPSC neuron, and primary CD8 T cell labeled with PKmito Orange (IMM), U2OS labeled with COX8a-mStayGold (IMM). The green cell in the bottom left corner in the primary T cell video is a Raji cell labeled with SybrGold and pre-treated with the superantigen SEE. All videos are taken on a SNOUTY microscope.

**Movie S3 (Movie03_classI_spontaneous_pearling.gif)**

Class I spontaneous mitochondrial pearling of primary human fibroblast characterized by calcium uptake and shrinking of the mitochondrial tubule. Cell labeled with COX8a-mStayGold (IMM, left-side) and Mito-R-GECO1 (mito-calcium, right-side). Video taken on SNOUTY microscope.

**Movie S4 (Movie04_iPSC_spon_action_potential.gif)**

Spontaneous pearling in iPSC-derived cells concurrent with action-potential firing. Neurons labeled with CalBryte^TM^ which marks cytosolic calcium (top-panel) and PKmito Orange (IMM, bottom-panel). All videos are taken on a SNOUTY microscope.

**Movie S5 (Movie05_iPSC_KCL_injection.gif)**

Action potential induced with 50mM KCl injection followed by mitochondrial pearling in iPSC-derived cells. Neurons labeled with CalBryte^TM^ which marks cytosolic calcium (left-panel) and PKmito Orange (IMM, right-panel). Video taken on SNOUTY microscope.

**Movie S6 (Movie06_Jurkat_Tcell_synapse.gif)**

Mitochondrial pearling during immunological synapse formation in Jurkat cells with Raji cell pre-treated with superantigen SEE. Jurkat cell labeled with COX8a-mStayGold (IMM) and SiR actin. Raji cell labeled with PKmito Orange (IMM). Video taken on SNOUTY microscope.

**Movie S7 (Movie07_Primary_CD8_Tcell_synapse.gif)**

Mitochondrial pearling at the synapse of activated primary human CD8⁺ T cells forming a synapse with an Raji cell pre-treated with superantigen SEE. CD8⁺ T cell labeled with PKmito Orange (IMM) and Raji cell labeled with SybrGold (mtDNA). Video taken on SNOUTY microscope.

**Movie S8 (Movie08_Tcell_synapse_calcium.gif)**

Calcium-flux–coupled with mitochondrial pearling at the T cell synapse. Jurkat cell labeled with COX8a-mStayGold (IMM) and Mito-R-GECO1 (mito-calcium) forming immune synapse with Raji cell labeled with SybrGold (mtDNA) and pre-treated with superantigen SEE.Video taken on SNOUTY microscope.

**Movie S9 (Movie09_senescenct_fibroblast_whole_cell_pearling.gif)**

Whole-cell mitochondrial pearling in primary human replicatively senescent fibroblast. Fibroblast has undergone 50 population doublings (∼50 divisions and 80 days of growth) and is now at minimal growth rate. Cell labeled with PKmito Orange (IMM). Video taken on SNOUTY microscope.

**Movie S10 (Movie10_U2OS_rotenone.gif)**

Mitochondrial pearling in U2OS cells treated with complex I inhibitor rotenone, 5 µM for 30min. Cell labeled with COX8a-mEmerald (IMM). Video imaged at 1 volume/second on SNOUTY.

**Movie S11 (Movie11_classII_spontaneous_pearling.gif)**

Class II spontaneous mitochondrial pearling in a primary human fibroblast characterized by elongation of the mitochondrial tubule without a rise in mitochondrial calcium. Cell labeled with COX8a-mStayGold (IMM, left-side) and Mito-R-GECO1 (mito-calcium, right-side). Video imaged at 1 volume/second on SNOUTY.

**Movie S12 (Movie12_calcium_rise.gif)**

Mitochondrial uptake in calcium coinciding with pearling in a U2OS cell. Cell labeled with Mito-R-GECO1 (mito-calcium) image on NIkon W1 spinning-disk confocal microscope.

**Movie S13 (Movie13_hypotonic_shock&recovery.gif)**

Hypotonic shock (0.1x PBS) induced cell-wide mitochondrial pearling and subsequent recovery to cylindrical morphology by return to isotonic media. Primary human fibroblast cell labeled with COX8a-mStayGold (IMM) imaged at 1 volume/second on SNOUTY.

**Movie S14 (Movie14_osmotic_shock_series.gif)**

Series of osmotic shocks demonstrating reversible pearling dynamics by swapping media between isotonic (DMEM) to hypotonic (0.1x PBS) to hypertonic (10x PBS) solutions. Primary human fibroblast labeled with PKmito Orange (IMM).

**Movie S15 (Movie15_hypertonic_recovery_pearling.gif)**

Hypertonic shock (10x PBS) followed by recovery-associated mitochondrial pearling in isotonic media. Cell labeled with COX8a-mStayGold (IMM, left-side) and Mito-R-GECO1 (mito-calcium, right-side) imaged at 1 volume/second on SNOUTY.

**Movie S16 (Movie16_chemical_inducers.gif)**

Mitochondrial pearling elicited by a different classes of chemical inducers including the calcium ionophore ionomycin (2µM), the potassium ionophore valinomycin (10µM), the BKCa channel activator NS1619 (10µM), the membrane permeabilization agent digitonin (10µM), and the chemical fixation agent paraformaldehyde (4%). All videos are mitochondria from different primary human fibroblasts labeled with COX8a-mStayGold (IMM) and taken at 1 volume/second on SNOUTY microscope.

**Movie S17 (Movie17_BKA_24fps.gif)**

Bongkrekic acid (BKA, 10 µM) injection marked by ‘hypertonic’ morphology preceding cell-wide mitochondrial pearling. Cell labeled with COX8a-mStayGold (IMM, left-side) and Mito-R-GECO1 (mito-calcium, right-side) imaged at 1 volume/second on SNOUTY.

**Movie S18 (Movie18_Digitonin_pearling.gif)**

Cell-wide mitochondrial pearling induced by injection of the membrane permeabilization agent digitonin (10 µM) in a primary human fibroblast labelled with COX8a-mStayGold (IMM) and imaged on SNOUTY at 1 volume/second.

**Movie S19 (Movie19_4PFA+0.05GA_fixation.gif)**

Mitochondrial pearling induced by chemical fixation with 4% PFA + 0.05% GA on mitochondrial morphology in a U2OS cell labelled with COX8a-mStayGold and imaged on SNOUTY at 1 volume/second.

**Movie S20 (Movie20_3PFA+2.0GA_fixation.gif)**

Mitochondrial morphology maintained after chemical fixation with 3% PFA + 2% GA on mitochondrial morphology in a primary human fibroblast cell labelled with COX8a-mStayGold and imaged on SNOUTY at 1 volume/second. Note, tubules appear partially ‘flaccid’ suggesting mild hypertonic effect of the fixative.

**Movie S21 (Movie21_ionomycin_hyperosmotic_rescue.gif)**

Hypertonic shock (1.5x PBS) rescuing cell-wide pearling induced by ionomycin injection (4 µM) on primary human fibroblast labeled with COX8a-mStayGold (IMM, left-panel) and Mito-R-GECO1 (mito-calcium, right panel). Video taken on SNOUTY at 1 volume/second.

**Movie S22 (Movie22_hypertonic_ionomycin_inhibition.gif)**

Cells pre-treated with hypertonic media (10x PBS) inhibiting ionomycin-induced (4 µM) pearling despite the rise in mitochondrial calcium. Pearling morphology appears after hypertonic media is replaced with isotonic media containing ionomycin (4 µM). U2OS cells labeled with COX8a-mStayGold (IMM, left-panel) and Mito-R-GECO1 (mito-calcium, right panel). Video taken on SNOUTY at 1 volume/second.

**Movie S23 (Movie23_long_laser_pearling.gif)**

Laser stimulation of mitochondrial tubule induced pearling across all connected tubules as far as 150 µm away from the point of stimulation. Video taken on nikon W1 widefield at 2 seconds/frame. Primary human fibroblast labelled with CellLight MitoGFP. The blue arrow indicates the point of stimulation by a focused 405nm laser fired at 20mW for 200 milliseconds with ∼0.33mW reaching the sample.

**Movie S24 (Movie24_whole_cell_laser_pearling.gif)**

Laser stimulation inducing mitochondrial pearling of the entire hyperfused mitochondria network in an untreated primary human fibroblast labeled with CellLight MitoGFP. The blue arrow indicates the point of stimulation by a focused laser fired at 20mW for 200 milliseconds with ∼0.33mW reaching the sample.

**Movie S25 (Movie25_laser_double_pearling.gif)**

A single laser stimulation induces successive pearling events around the point of stimulation. The blue arrow indicates the point of stimulation by a focused laser. Untreated primary human fibroblast labeled with CellLight MitoGFP. The blue arrow indicates the point of stimulation by a focused 405nm laser fired at 20mW for 200 milliseconds with ∼0.33mW reaching the sample.

**Movie S26 (Movie26_calcium_preceding_laser_pearling.gif)**

Mitochondrial calcium transiently increased preceding laser-induced pearling. The blue arrow indicates the point of stimulation by a focused 405nm laser fired at 20mW for 200 milliseconds with ∼0.33mW reaching the sample. Untreated primary human fibroblast labeled with COX8a-mStayGold (IMM, left-panel) and Mito-R-GECO1 (mito-calcium, right-panel).

**Movie S27 (Movie27_FCCP_laser_pearling.gif)**

Inhibition of laser induced pearling by depolarizing mitochondria with the proton ionophore FCCP (10 µM for 25 min) treated on U2OS cells labeled with COX8a-mEmerald (IMM). The blue arrows indicate the points of stimulation by a focused 405nm laser fired at 20mW for 200 milliseconds with ∼0.33mW reaching the sample.

**Movie S28 (Movie28_Actin_inhibitors.gif)**

Impact of actin-polymerization inhibitors on mitochondrial pearling recovery after laser stimulation. Cells treated with latrunculinB (1 µM for 1 hour), CK666 (85 µM for 1 hour) and y27632 (20 µM for 1 hour) and labelled with MitoTracker Green. The blue arrows indicate the points of stimulation by a focused 405nm laser.

**Movie S29 (Movie29_TMRE_collapse&recovery.gif)**

Mitochondrial membrane potential tracked with TMRE fluorescence showing membrane-potential collapse and recovery after laser-induced pearling. Primary human fibroblast labeled with CellLight MitoGFP (mito-matrix, left-panel) and TMRE (MMM, middle-panel). The blue arrows indicate the points of stimulation by a focused 405nm laser fired at 20mW for 200 milliseconds with ∼0.33mW reaching the sample.

**Movie S30 (Movie30_STED_cristae.gif)**

Live-cell STED super-resolution imaging of mitochondrial cristae reorganization during pearling by stimulation with a 405nm laser. Primary human fibroblasts labeled with PKmito Orange. The blue arrow indicates the point of stimulation by a focused 405nm laser fired at 20mW for 200 milliseconds. Single non-STED confocal frame shown right after the point of stimulation before returning to STED imaging.

**Movie S31 (Movie31_ER_dependent_pearling.gif)**

Spontaneous mitochondrial pearling with endoplasmic reticulum network wrapped around pearl structure. U2OS cells labeled with COX8a-mEmerald (IMM) and CellLight-ER-RFP (calreticulin and KDEL).

**Movie S32 (Movie32_ER_independent_pearling.gif)**

Spontaneous mitochondrial pearling form without direct overlap of pearl constriction points with ER. U2OS cells labeled with COX8a-mEmerald (IMM, left-panel) and CellLight-ER-RFP (calreticulin and KDEL, middle-panel).

**Movie S33 (Movie33_ER_independent_pearling_v2.gif)**

Spontaneous mitochondrial pearling form without overlap of pearl constriction points with ER. U2OS cells labeled with COX8a-mEmerald (IMM, left-panel) and CellLight-ER-RFP (calreticulin and KDEL, middle-panel).

**Movie S34 (Movie34_microneedle_control_pearling.gif)**

Microneedle manipulation of untreated U2OS cell inducing pearling on selected mitochondrial tubule by mechanical force. The needle first is pushed against the mitochondrion, removed from the cell and then re-applied for second manipulation before the cell itself begins to retract from cover slip. Blue dot indicates the tip of the needle marked by fluorescent coating. U2OS cell labeled with COX8a-mStayGold (IMM, left-panel) and Mito-R-GECO1 (mito-calcium, right-panel).

**Movie S35 (Movie35_microneedle_FCCP_pearling.gif)**

Microneedle manipulation of mitochondria in U2OS cells depolarized with the proton ionophore 10 µM FCCP (30min). The needle fails to induce pearling and fragments the mitochondrial tubule at the point of contact. Blue dot indicates the tip of the needle marked by fluorescent coating. U2OS cell labeled with COX8a-mStayGold (IMM, left-panel) and Mito-R-GECO1 (mito-calcium, right-panel).

**Movie S36 (Movie36_microneedle_puncture_pearling.gif)**

Microneedle manipulation of mitochondria in unrtreatd U2OS cells causign cell wid-pearling and retraction of the cell from the coverslip. Blue dot indicates the tip of the needle marked by fluorescent coating. U2OS cell labeled with COX8a-mStayGold (IMM, left-panel) and Mito-R-GECO1 (mito-calcium, right-panel).

**Movie S37 (Movie37_microneedle_latB_pearling_with_calcium.gif)**

Microneedle manipulation of mitochondria in U2OS cells with actin depolymerized with 1 µM latrunculin B (30 min). The needle induces pearling specifically on the portion of mitochondria undergoing mechanical stretching without nearby tubules changing. Blue dot indicates the tip of the needle marked by fluorescent coating. U2OS cell labeled with COX8a-mStayGold (IMM, left-panel) and Mito-R-GECO1 (mito-calcium, right-panel).

**Movie S38 (Movie38_microneedle_latB_pearling_without_calcium.gif)**

Microneedle manipulation of mitochondria in U2OS cells with actin depolymerized with 1 µM latrunculin B (30 min). The needle induces pearling without a simultaneous rise in mitochondrial calcium. Blue dot indicates the tip of the needle marked by fluorescent coating. U2OS cell labeled with COX8a-mStayGold (IMM, left-panel) and Mito-R-GECO1 (mito-calcium, right-panel).

